# Smc5/6 - microtubules binding shapes pericentromeric chromatin

**DOI:** 10.1101/2024.11.13.623393

**Authors:** Ànnia Carré-Simon, Cecilia Martuzzi, Sarah Isler, Renaud Batrin, Henrik Dahl Pinholt, Timothy Földes, Guillaume Laflamme, Maria Barbi, Leonid Mirny, Damien D’Amours, Emmanuelle Fabre

**Affiliations:** Université de Paris Cité, INSERM, CNRS, IRSL, U1342, EMR8000, Paris, France; Ottawa Institute of Systems Biology, Department of Cellular and Molecular Medicine, Ottawa, Canada; Department of Physics, Massachusetts Institute of Technology, Cambridge, MA, USA; Sorbonne Université, CNRS, LPTMC, Paris, France; Institute for Medical Engineering and Science, Massachusetts Institute of Technology, Cambridge, MA, USA; Department of Pathology, NYU Grossman School of Medicine, New York, NY, USA

**Keywords:** Smc5/6, microtubules, pericentromeres, mitotic spindle, DNA double-strand break repair, chromatin dynamics

## Abstract

Centromeres and pericentromeres are specialized chromatin regions essential for accurate chromosome segregation. Smc5/6, which localizes at pericentromeres, can bind microtubules, yet its role in chromatin folding is unclear. Here, we investigate the functional relevance of Smc5/6- microtubule binding in yeast, by targeting two lysines (K624, K631) within the Smc5 hinge domain known to mediate this binding. Using high-temporal-resolution imaging, polymer modelling, and *in vitro* approaches with a separation-of-function mutant *smc5-2KE*, we demonstrate that microtubules binding by Smc5/6 constrains chromatin dynamics and promotes pericentromeric folding. The *smc5-2KE* mutant, combined with a hypomorphic kinetochore mutant (Mtw1-3xGFP), leads to spindle and cytokinesis defects and triggers the spindle checkpoint. Furthermore, homologous recombination repair in pericentromeres is compromised. Overall, our findings indicate that Smc5/6 – microtubules association safeguard pericentromeric architecture and genome stability during mitosis.

## Introduction

The three-dimensional architecture of the genome governs gene expression, DNA replication and repair (Misteli, 2020). Centromeres are specialized regions within this chromosomal architecture that serve as scaffolds for microtubule attachment via the kinetochore, a complex of around 80 proteins (Biggins, 2013; Talbert and Henikoff, 2020). Surrounding the centromere, pericentromeric regions adopt a distinct chromatin configuration that is essential for genome stability (Bloom, 2024). Pericentromeric configuration and centromere/kinetochore assembly are crucial to counterbalance the mechanical forces exerted by spindle microtubules during mitosis. Yet, how pericentromeric structure is shaped to resist the forces exerted by microtubules is poorly understood (Harasymiw et al., 2019; Stephens et al., 2011). In budding yeast, pericentromeres span around 30-50 kb (Paldi et al., 2020) and are highly enriched in structural chromosome maintenance (SMC) complexes, which include cohesins, condensins and the Smc5/6 complex (Smc5/6 in short) (D’Ambrosio et al., 2008; Jeppsson et al., 2014; Megee et al., 1999; Tanaka et al., 1999). Cohesins and condensins participate in the establishment of DNA loops that resist mitotic spindle forces (Lawrimore and Bloom, 2022). While assembly of an Smc5/6 dimer was recently shown to contribute to DNA loop folding *in vitro* (Pradhan et al., 2023), the folding function of Smc5/6 at pericentromeres is unknown.

Smc5/6 is composed of Smc5, Smc6 and non-SMC proteins (Nse1, Mms21 and Nse3-6) (Roy et al., 2024). Smc5 and Smc6 consist of antiparallel coiled-coil arms that dimerize via a hinge domain at one end and are linked at the other end by their ATPase head domains. As a key player in genome stability that mediates post-translational modifications (PTMs), Smc5/6 interacts with many chromosome proteins during its cellular functions (Roy et al., 2024). One of the most intriguing interactors of the complex is the alpha/beta tubulin dimer. We previously showed *in vitro* that the hinge domain of the Smc5 protein binds the tubulin dimer exclusively in the form of microtubule polymers. The hinge domain contains three lysines (K624, K631 and K667) that interact directly *in vitro* with the C-terminal negatively charged tails of tubulins (Laflamme et al., 2014). A recent mass spectrometry analysis of overexpressed Smc5/6 purified from exponentially growing yeast, which we have confirmed, also identified alpha/beta tubulin subunits Tub1 and Tub2, as cofactors binding to the Smc5/6 complex (Gutierrez-Escribano et al., 2020, Figure supplement 1A-C). These data support an interaction between the Smc5/6 complex and microtubules, but the function of such a link remains an open question.

Besides its enrichment at pericentromeres, Smc5/6 is also found at telomeres and rDNA to maintain their organization (Moradi-Fard et al., 2016; Torres-Rosell et al., 2005; Zhao and Blobel, 2005). For example, deletion of the Nse2/Mms21 subunit of the Smc5/6 leads to defects in genome organization, including telomere declustering and nucleolar fragmentation (Moradi-Fard et al., 2016; Torres-Rosell et al., 2005; Zhao and Blobel, 2005). Additionally, smc5/6 mutants (*smc6-9*) exhibit chromosome segregation defects, notably in repetitive regions such as rDNA and telomeres (Torres-Rosell et al., 2007, 2005). Beyond its structural role, Smc5/6 participates in genome integrity by promoting the repair of damaged DNA(Aragón, 2018; Roy et al., 2024). Smc5/6 is recruited within ∼5kb of double strand breaks (DSB (De Piccoli et al., 2006; Lindroos et al., 2006), holds the DSB ends together (Phipps et al., 2024) and facilitates homologous recombination (HR) (Ampatzidou et al., 2006; Bermúdez-López et al., 2010; Pebernard et al., 2006; Torres-Rosell et al., 2005), a DNA repair process that uses a homologous sequence as a template to accurately repair DSBs (Symington and Gautier, 2011).

Here, we examine the role that the association between Smc5/6 and microtubules plays in pericentromeric chromatin folding. We combine mutational analysis and fluorescence microscopy in living cells with polymer simulations to show that the association between microtubules and Smc5/6 impacts pericentromeric chromatin folding, dynamics and integrity.

## Results

### The *smc5-2KE* mutant leads to increased chromatin dynamics at pericentromeres

To determine how Smc5/6 and microtubules could interact, we analyzed the spatial distribution of K624, K631 and K667 residues using USCF Chimera based on the hinge domain (PDB:7YLM (Li et al., 2024)). Our structural assessment revealed that K624 and K631 are positioned on the outer surface of the hinge domain, making them accessible for potential interactions with microtubules. In contrast, K667 is embedded in the coiled-coil arm of Smc5 and could not theoretically interact with microtubules (**Figure 1A**). These results agree with the position of K624 and K631 in the specific loop of the C domain unique to the Smc5/6 according to the X-ray crystal structures of the *S. pombe* Smc5 hinge domain (Alt et al., 2017).

**Figure 1:**
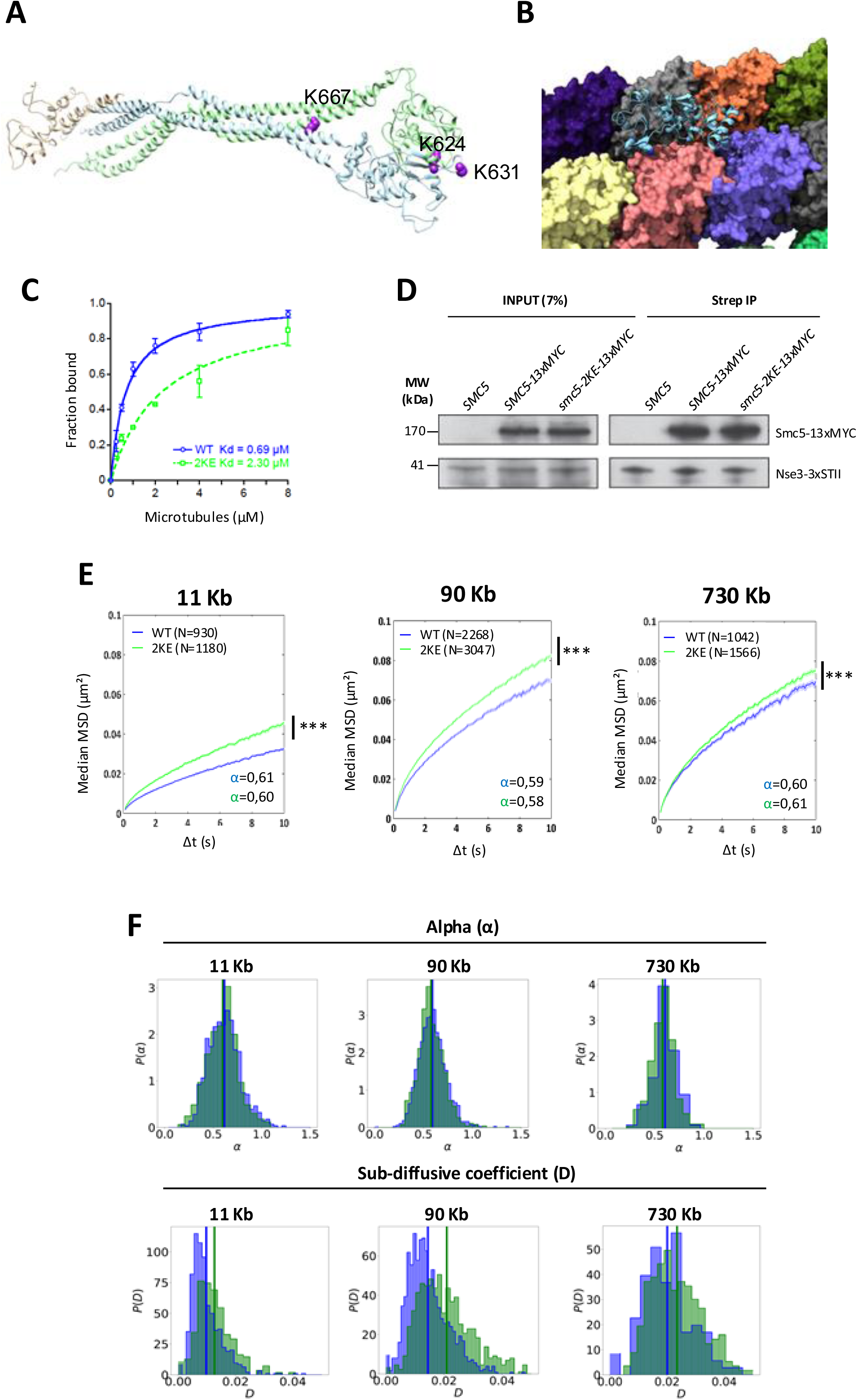
The *smc5-2KE* mutant shows decreased binding affinity to microtubules *in vitro* and leads to an increased chromatin dynamics at pericentromeres and beyond. (A) Representation of the CryoEM of the Smc5-Smc6 hinge from Li et al., NSMB, 2024 (PDB: 7YLM). The Smc5 hinge domain is shown in blue, the Smc6 hinge in green, and Nse2 in brown. Lysines 624, 631, and 667 are highlighted in purple. **(B)** AlphaFold model of the Smc5 hinge (amino acids 450–645) showing a high predicted probability of interaction with tubulin dimers. Lysines 631 and 624 (in purple) contact adjacent tubulin subunits in the microtubule. Smc5 is shown as a cartoon in light blue; tubulin dimers are colored individually. **(C)** The fraction of microtubule-bound Smc5 and Smc5-2KE from 3 independent experiments was plotted as a function of microtubules (MT) concentrations. Dissociation constants (Kd) were determined by curve fitting to a hyperbola. **(D)** Co-immunoprecipitation (co-IP) experiments showing the association of Nse3-3xSTII with Smc5 or Smc5-2KE proteins (both tagged with C-terminal 13xMYC epitopes) in yeast extracts. A whole cell extract was loaded as an input (7%; **left**) and Smc5/6 complex components were immunoprecipitated from this extract using StrepTactin XT® resin (**right**). For each panel, the first lane represents a negative control where only Nse3 was tagged (with 3xSTII), whereas the second and third lanes show Nse3-3xSTII co-purified with Smc5-13xMYC and Smc5-2KE-13xMYC proteins, respectively. **(E)** Mean square displacement (MSDs) as a function of increasing time intervals in the WT (blue) and in the *smc5*-2KE mutant (green) for a locus at different positions of chromosome IV (11kb, 90 kb and 730 kb, from left to right). Wilcoxon rank-sum test results between distributions, with the p-value. * (p<0.05), *** (p<0.0001). **(F)** Distributions of the exponent alpha (upper line) and of the sub-diffusion coefficient D (lower line) for the WT (blue) and in the *smc5-2KE* mutant (green) for loci at different positions of chromosome IV. The values of parameters alpha and D are calculated from single trajectories as described in Material and Methods. Vertical lines correspond to average values (WT in blue, *smc5-2KE* mutant in green).

In a second step, using AlphaFold3 to model alpha/beta tubulin subunits and a segment of the Smc5 hinge domain containing lysines K624 and K631 we identified a high probability of interaction between this region and microtubules (iPTM = 0.72) (**Figure 1B**), in contrast with the low probability of interaction with other proteins such as actin (iPTM=0.16) or the subunit Shs1 of the septin complex (iPTM=0.18) (**Figure supplement 2A**).

This observation led us to focus on K624 and K631 and modify their electrostatic attraction with the C-terminal tubulin tails by mutating them to negatively charged glutamic acid residues, generating the *smc5-2KE* mutant (Material and methods and Laflamme et al., 2014). The microtubule-binding capacity of the *smc5-2KE* mutant was verified *in vitro* by comparing the co-sedimentation of purified Smc5 and Smc5-2KE proteins with microtubules. We observed a 3-fold reduction in the binding affinity of Smc5-2KE compared to wild-type (WT) Smc5 (**Figure 1 C – Figure supplement 2B**). The reduced affinity of Smc5-2KE for microtubules was not due to a defect in the mutant’s interaction with other proteins in the Smc5/6 complex, as indicated by co-immunoprecipitation (CoIP) of the Nse3 subunit with both WT Smc5 and the Smc5-2KE (**Figure 1D**). The preservation of Smc5/6 function in the *smc5-2KE* mutant is consistent with the absence of significant defects in cell growth, cell division, cell cycle progression, sensitivity to DNA damage agents or thermosensitivity, as demonstrated in mutated cells (**Figure supplement 2C-H**). Furthermore, the *smc5-2KE* mutant did not impact the binding with DNA, as observed by ChIP-qPCR at centromeres, telomeres and rDNA (**Figure supplement 2I**), indicating that the associations of Smc5/6 with microtubules and with chromatin are independent. Therefore, we used *smc5-2KE* as a separation-of-function mutant that is specifically defective in microtubule binding, to explore the role of Smc5/6 association to microtubules in chromatin folding and dynamics *in vivo*.

To first investigate the impact of weakened microtubule-Smc5/6 association on chromatin dynamics as a function of chromosome position, we tracked a GFP-labelled chromatin array (tetO-TetR-GFP) positioned at 11, 90, and 730 kb from centromere IV (CEN IV) in both WT and *smc5-2KE* mutant. We calculated the mean square displacement (MSD) as previously described and extracted different dynamic parameters (Cabal et al., 2006; García Fernández et al., 2021; Hajjoul et al., 2013; Herbert et al., 2017; Marshall et al., 1997; Miné-Hattab et al., 2017; Miné-Hattab and Rothstein, 2012) (Material and methods). In WT cells, the different tracked loci followed a sub-diffusive movement characterized by an anomalous *α* exponent below 1.0 (between 0.59 and 0.61, **Figure 1E - Figure supplement 3A**). These values of *α*, typical of self-avoiding bead-spring polymer models (Panja and Barkema, 2009), indicate that the connectivity of the chromosome, qualitatively determining the nature of its dynamics, is independent of its distance from the centromere. However, the sub-diffusive coefficient D, as well as MSD at 10s (MSD_10s_), indicated higher mobility at positions further from the centromeric attachment point to kinetochore microtubules on CEN IV, as previously documented (**Figure 1E,F - Figure supplement 3A-C**) (Cabal et al., 2006; Hajjoul et al., 2013; Herbert et al., 2017; Neumann et al., 2012; Verdaasdonk et al., 2013). Interestingly, sub-diffusive chromatin motion was maintained in *smc5-2KE* mutated cells with a similar *α* parameter, but chromatin dynamics increased, characterized by an increase in D, at all loci compared to WT cells. An increase in D can be explained by a decrease of interactions within the chromatin polymer or with external factors, consistent with the loss of centromeric constraint and/or partial detachment of centromeric chromatin from kinetochore microtubules (Socol et al., 2019). Of note, the increase in chromatin dynamics in mutated cells was more pronounced the closer the locus was to the centromere (**Figure 1E,F**). MSD_10s_ increased in the *smc5-2KE* mutant, compared to WT, by factors of 1.36, 1.18 and 1.07 for distance 11, 90 and 730 kb from CEN IV, respectively (**Figure supplement 3C**). Increased chromatin mobility of the *smc5-2KE* mutant was verified for another locus located at 90 kb from CEN V (**Figure supplement 3D**) and observed at different phases of the cell cycle, including G1 and metaphase (**Figure supplement 4**).

Since lysine residues undergo a wide range of reversible PTMs such as SUMOylation, acetylation and ubiquitination, which can regulate protein functions, including protein-protein interactions (Wang and Cole, 2020), we mutated both lysines to arginine (a positively charged amino acid unaffected by PTM), to determine whether lysine PTMs are involved in the Smc5 effect on chromatin mobility. The *smc5-2KR* mutant showed chromatin mobility like WT (**Figure supplement 3D**) indicating that the positive charges at amino-acid positions 624 and 631 of Smc5, rather than their potential to be modified by PTMs, are required to promote normal chromatin dynamics *in vivo*.

Finally, since sister chromatid cohesion has been shown to restrict chromatin dynamics during S phase (Cheblal et al., 2020; Dion et al., 2013), we investigated whether cohesion defects in the *smc5-2KE* mutant could account for the observed changes in chromatin dynamics. To do so, we analysed TetO-TetR-GFP at 11 kb from CENIV in mitotically synchronized cells. We analysed the state of sister chromatid cohesion by quantifying the number of cells displaying either one GFP focus (cohesive chromatin) or two GFP foci (defective cohesion) (Garcia-Luis et al., 2022; Paldi et al., 2020). Our analysis revealed no significant differences in the number of cells with single GFP foci between the *smc5-2KE* mutant and the WT strain (**Figure supplement 2E**), indicating no cohesion defects. In contrast, the deletion of the Smc1 cohesin subunit, used as a control for defective cohesion, resulted in the separation of GFP foci (**Figure supplement 2F**) (Garcia-Luis et al., 2022).

Altogether, our results support the hypothesis that lysines K624 and K631, conserved in the C domain of Smc5 hinge, limit chromatin dynamics possibly through the attraction of their positive charge to microtubules, as observed *in vitro*.

### Increased chromatin dynamics of the *smc5-2KE* mutant is not linked to spontaneous DNA damage or loss of Smc5/6

Increase in chromatin dynamics is part of the DNA Damage Response (García Fernández et al., 2022b; Hauer and Gasser, 2017). To determine whether increased chromatin mobility of the *smc5-2KE* mutant results from spontaneous DNA damage, we first monitored Rad52 foci, as a marker of DNA repair at sites of damage (Lisby et al., 2001). While treatment with the genotoxic agent Zeocin yielded a similarly high number of Rad52 foci in WT and *smc5-2KE* mutant (74% ± 6.7 and 79,5% ± 7.8 foci respectively), we observed comparable low levels of Rad52 foci in WT and *smc5-2KE* strains in the absence of Zeocin (7% ± 1.1 and 8.9% ± 1.3 foci respectively) (**Figure supplement 5A**). We also analyzed Rad53 phosphorylation, a checkpoint modification induced by DNA damage (Pellicioli et al., 1999). Rad53 phosphorylation was not detected in *smc5-2KE* cells in the absence of Zeocin, in contrast to Zeocin-treated WT and mutant cells (**Figure supplement 5B**). Finally, we investigated the phosphorylation of histone H2A (γH2A), a conserved PTM induced by DNA damage (Shroff et al., 2004). Upon Zeocin treatment, WT and *smc5-2KE* cells displayed increased levels of γH2A, while γH2A background levels were similar in untreated strains (**Figure supplement 5C**). These observations are consistent with the lack of Rad53 phosphorylation and γH2A observed upon Smc5/6 complex degradation (Peng et al., 2018). Taken together these results establish that the increased chromatin dynamics observed in the *smc5-2KE* mutant are not due to higher levels of spontaneous DNA damage.

K624E and K631E mutations do not affect the stability of Smc5/6 *in vitro* (**Figure 1D**), but we could not exclude that a defect in the Smc5/6 complex, undetectable *in vitro*, could account for the increased chromatin mobility of the *smc5-2KE* mutant *in vivo*. To address this point, we investigated chromatin mobility in cells defective for Smc5/6 by specifically degrading Smc5 or Smc6 proteins using the auxin-inducible degron (AID) strategy. Cells expressing AID-tagged Smc5 or Smc6 proteins grew like WT cells in the absence of auxin but failed to grow in the presence of 1 mM auxin (**Figure supplement 6A-B**). This indicates efficient degradation of the tagged proteins and confirms that complete loss of Smc5 and Smc6 proteins is incompatible with cell viability, unlike the separation-of-function phenotype of the *smc5-2KE* mutant. Degradation took place 30 minutes after auxin addition, as shown by western blot for Smc5-AID (**Figure supplement 6C**). Surprisingly, a reduction in chromatin dynamics in the pericentromeric region, at 11 kb from the centromere, was observed when Smc5-AID or Smc6-AID protein degradation was induced by 1h auxin addition (**Figure supplement 6D,F**). These findings contrast with the increase in chromatin dynamics we observed upon degradation of the Smc1 subunit of cohesin (**Figure supplement 6E,F**), as previously documented (Cheblal et al., 2020; Verdaasdonk et al., 2013). The opposite effects of Smc5/6 and cohesins on chromatin mobility suggest different mechanisms, which in the case of cohesins might depend on their loop extrusion activity (Costantino et al., 2020; Dauban et al., 2020). Taken together, these results establish that Smc5/6 is required in maintaining chromatin dynamics which is itself constrained by the association of Smc5 with microtubules.

### The binding of microtubules to kinetochores is necessary but not sufficient to control pericentromeric chromatin dynamics

The constraint of pericentromeric chromatin dynamics is in line with studies in budding yeast, showing that release of centromeres from their attachment to the SPB upon microtubule depolymerization or centromere inactivation leads to enhanced chromatin mobility (Amitai et al., 2017; Marshall et al., 1997; Strecker et al., 2016; Verdaasdonk et al., 2013). We therefore investigated chromatin mobility following microtubule depolymerization by nocodazole in WT and in the *smc5-2KE* mutant. We exposed cells to a short treatment (30 minutes) with nocodazole (**Figure 2A**) and analyzed chromatin dynamics at different chromosome locations. This treatment depolymerized microtubules, as confirmed by the absence of microtubule filaments visualized by Tub2-GFP (**Figure 2B**) but did not block cycling of the cells (**Figure 2C**). In WT cells, chromatin mobility increased upon exposure to nocodazole at loci far from the centromere, *i.e.* at 90 kb and 730 kb from the centromere, as previously observed (**Figure 2D,G**) (Amitai et al., 2017; Marshall et al., 1997). In contrast, in the *smc5-2KE* mutant, nocodazole had no significant effect on chromatin mobility, indicating that the lack of microtubules or the Smc5-2KE mutation leads to a similar increase in chromosome dynamics with no cumulative effect of the two. Close to the centromere at 11kb from CEN IV, the increase in chromatin dynamics caused by nocodazole nevertheless reached a lower level than that observed in the *smc5-2KE* mutant (MSD_10s_ compared to WT, 1.14 vs. 1.32 respectively, **Figure 2D**). These observations support the view that binding of microtubules to Smc5/6 limits chromatin mobility throughout the chromosome and that in the pericentromeric region, Smc5/6 plays an additional role, possibly affecting chromatin properties.

**Figure 2:**
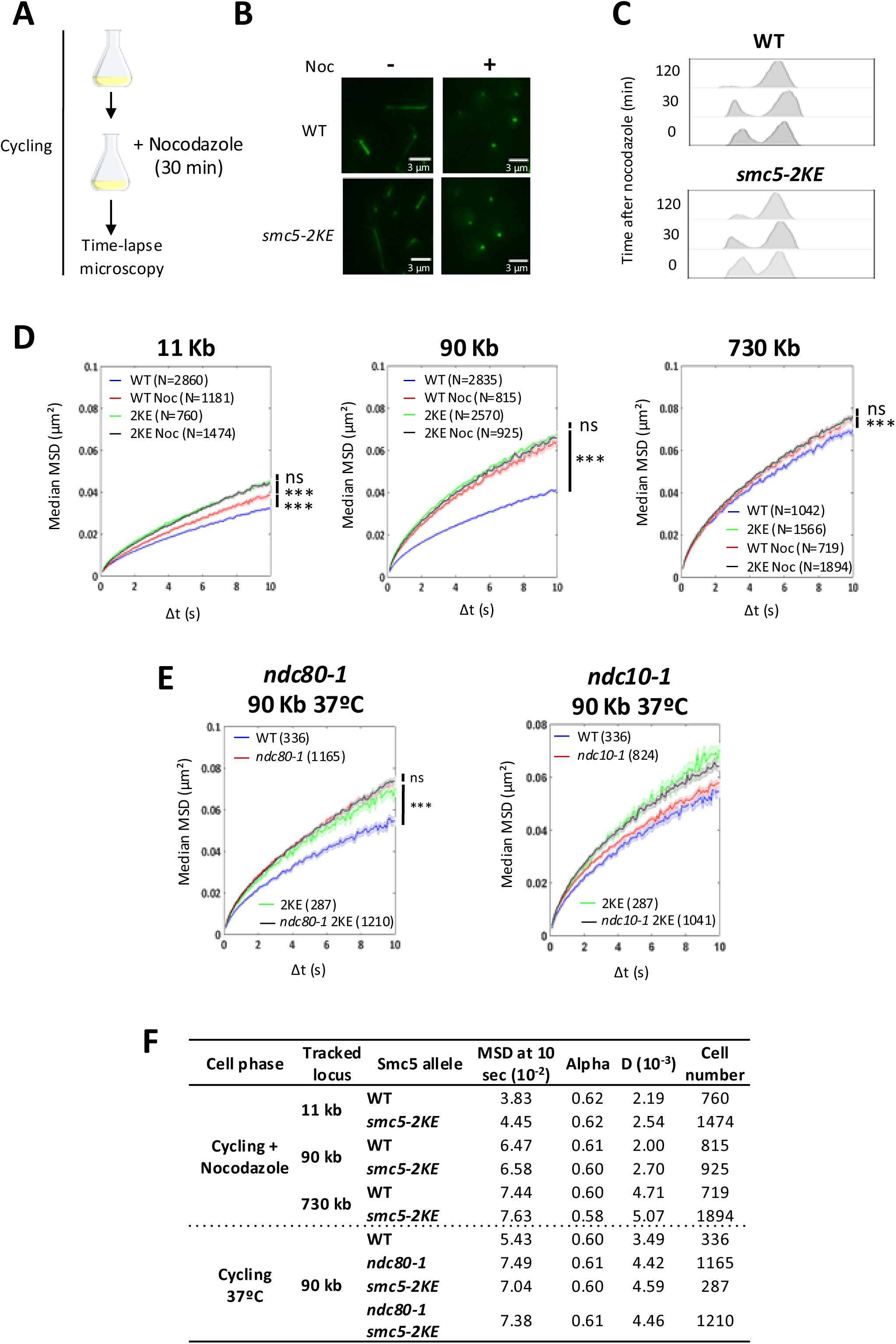
The binding of microtubules to kinetochore is necessary but not sufficient to control pericentromeric chromatin dynamics. **(A)** Experimental protocol used to depolymerize microtubules with 15 ng/µl of nocodazole while continued cycling. **(B)** Tub2-GFP in the WT and in the *smc5-2KE* mutant with or without addition of nocodazole. **(C)** Cells synchronization monitored by flow cytometry in cells treated with nocodazole (15 ng/µl). Samples were taken at 0, 30 and 120 minutes after nocodazole addition. **(D)** MSD as a function of increasing time intervals in the WT and *smc5*-2KE mutant at 11, 90 and 730 kb with or without the presence of nocodazole. Wilcoxon rank-sum test results between distributions, with the p-value “ns” non-significant, *** (p<0.0001). **(E)** MSDs as a function of increasing time intervals of the locus at 90 kb in the *ndc80-1* mutant (**left**) and *ndc10-1* mutant (**right**) combined with or without the Smc5-2KE mutation at 37°C. Wilcoxon rank-sum test results between distributions, with the p-value “ns” non-significant, *** (p<0.0001) **(F)** Table summarizing the different conditions with values extracted from the MSD: MSD at 10 s, the anomalous exponent (alpha), the sub-diffusive coefficient (D) and the number of cells analyzed.

In a complementary approach, we analyzed chromatin dynamics in two kinetochore thermosensitive mutants *ndc80*-1, which impairs kinetochores-microtubules binding (Wigge and Kilmartin, 2001) and *ndc10*-1, which affects centromeric chromatin binding (Lechner and Carbon, 1991). To effectively compare these mutants to *smc5-2KE*, we first confirmed that the chromatin behavior reported in the *smc5-2KE* mutant at 30°C is also observed at 37°C, the restrictive temperature for these mutants. Chromatin dynamics at 90 kb from CEN V displayed a comparable increase between *ndc80-1* and *smc5-2KE* single mutants at 37°C (**Figure 2E left, F**), with no additive effect in the *ndc80-1 smc5-2KE* double mutant (**Figure 2E left, F**). Conversely, the *ndc10-1* mutant showed no increase in chromatin dynamics compared to the WT, and an increase in chromatin mobility when combined with the Smc5-2KE mutation (**Figure 2E right, F**). These results indicate that only kinetochore mutations disrupting microtubule binding and not chromatin binding have an impact on chromatin dynamics.

Our analysis collectively demonstrates that changes in microtubule binding, whether due to KE mutations in the hinge of Smc5, kinetochore mutants that impair binding with microtubules or microtubule depolymerization, are sufficient to enhance chromatin dynamics away from the centromere. They also indicate that Smc5/6 plays an additional role controlling pericentromeric dynamics through its binding to microtubules.

### The *smc5-2KE* mutant modifies pericentromeric chromatin properties

The results above suggested that *smc5-2KE* mutant may impact chromatin properties at pericentromeres. We therefore compared intra-chromosomal distances of different loci in WT and mutant cells. Changes in intra-chromosomal distances underline changes in compaction, rigidity or loop extrusion activity (Hauer et al., 2017; Herbert et al., 2017; Phipps et al., 2024). We used strains R1 and R3 where pairs of differently fluorescently labelled loci, distant by ∼180 kb, are scattered on chromosome IV (Herbert et al., 2017) (Material and methods). The R1 strain has two labelled loci, one in in a pericentromeric position (at 4 kb from CEN IV) and one outside (**Figure 3A**) and the R3 strain has two labelled loci located centrally on the chromosome arm (at 735 kb from CEN IV, **Figure 3B**). The *smc5-2KE* mutation was introduced in both strains, and 2D distances between pairs of loci were measured from 3D images (Material and methods). As a control, we used Zeocin, which promotes an increase in intra-chromosomal distances at all positions, as previously shown (**Figure 3A,B**) (Hauer et al., 2017; Herbert et al., 2017). While intra-chromosomal distances between pair of loci in R3 strain remained similar between R3 WT and R3 *smc5-2KE* strains (**Figure 3B**), a significant increase in intra-chromosomal distances was observed in the R1 *smc5-2KE* mutant compared to the R1 WT strain (**Figure 3A**). This increase suggests that at this sub-chromosomal scale, changes in chromatin properties occur in the *smc5-2KE* mutant, which are revealed when at least one locus is pericentromeric. Although these data do not precisely define whether chromatin decompaction or stiffening are induced by the *smc5-2KE* mutant, they do indicate that the K624 and K631 dependent association of Smc5 with microtubules is required to establish or maintain pericentromeric chromatin.

**Figure 3:**
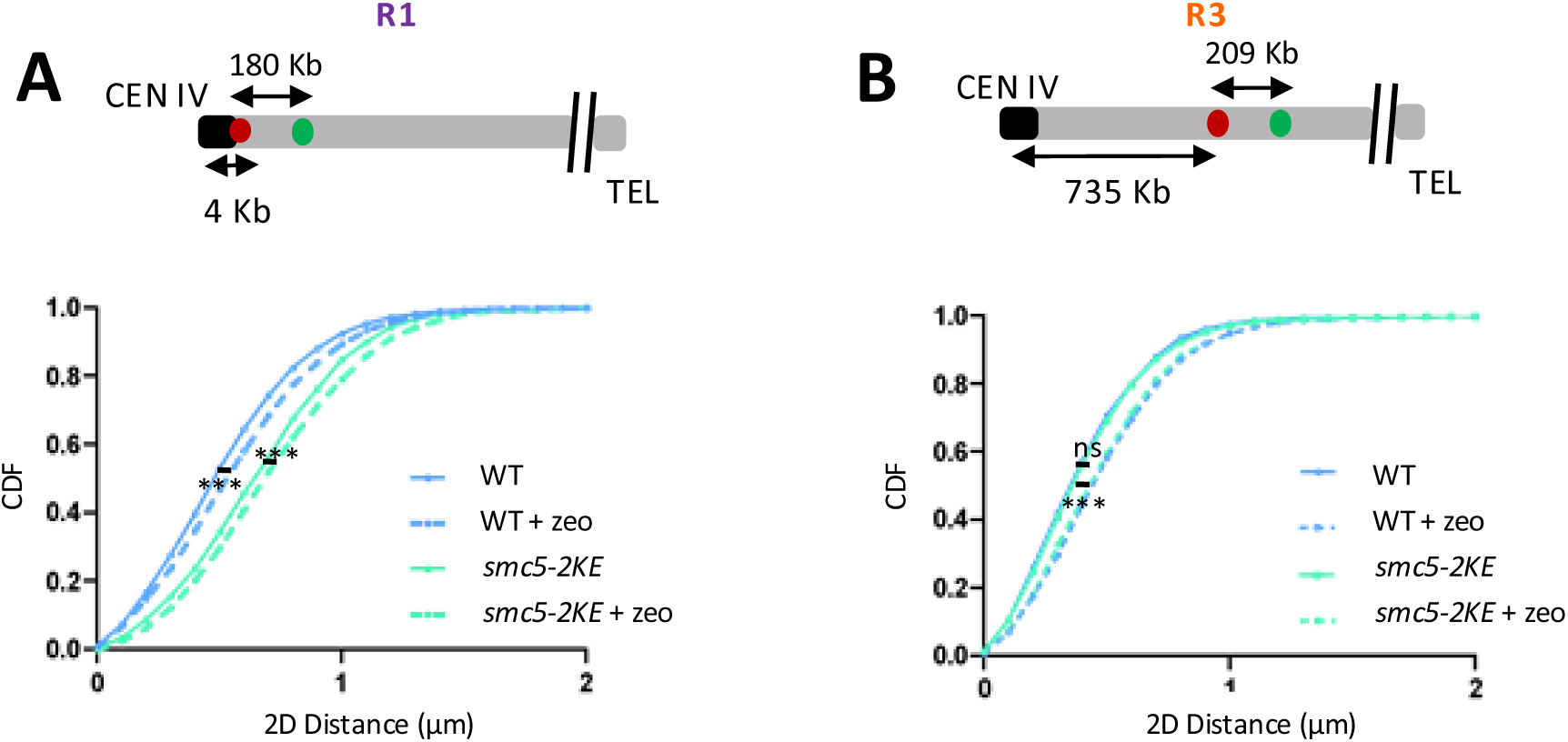
The *smc5-2KE* mutant changes chromatin properties in the pericentromeric region. (A-B) Cumulative distribution functions (CDF) show the intrachromosomal distances distribution measured for two pairs of loci 180 kb apart at two different positions of the chromosome IV, **(A)** close to the centromere (R1) and **(B)** in the middle of the arm (R3) in the presence or absence of Zeocin. Three experiments were done in each case: R1: N_WT_=21806, N_2KE_=9014, N_WT ZEO_=4091, N_2KE ZEO_=1570 and for R3: N_WT_=2778, N_2KE_=4403. Wilcoxon rank-sum test was used to analyze with *** for P<0.001.

### Polymer model recapitulates changes in chromatin dynamics at pericentromeres

We next used biophysical polymer simulations (Material and methods) to determine if in the pericentromeric region, microtubule binding and chromatin structure together could drive chromatin dynamics. Initially, we assessed chromatin mobility at positions corresponding to 11, 23, 90 and 730 kb from the centromere of different chromosomes. Polymer simulation showed that the increase of MSD_10s_ was proportional to the distance to the centromere as observed *in vivo* and plateaued at 90 kb (**Figure 4A** compared to **Figure 1C**). Although the increase in mobility in the polymer model was less pronounced than in the experimental data, the relative features were present and would allow investigation of the effects of perturbations.

**Figure 4:**
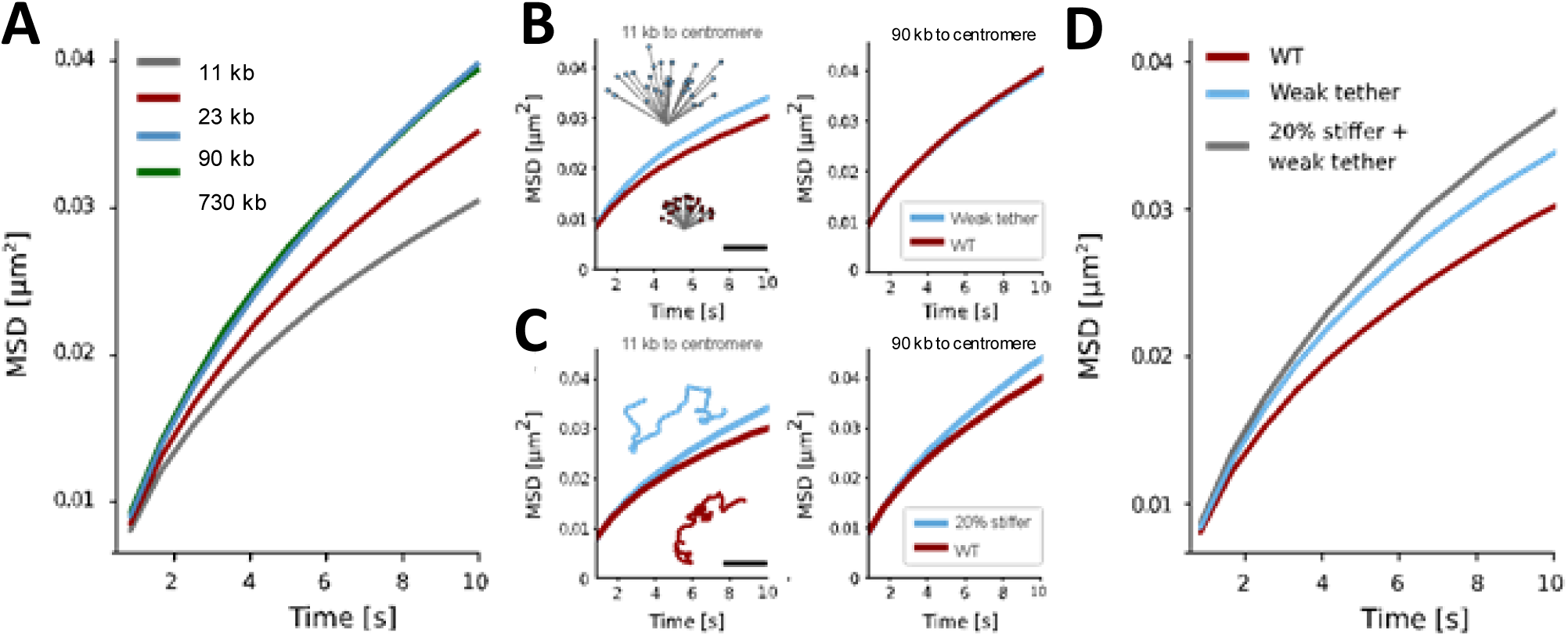
Polymer model recapitulates changes in chromatin dynamics at pericentromeres. **(A)** Comparison of MSDs for loci with various distances to the centromere. **(B)** Effect of increasing the flexibility of the SPB tether from rigid (red curves) to a spring constant of 1kbT/ (105 nm)^2^ which would allow for a standard deviation of 74 nm (105 nm/sqrt(2)) if no other forces were applied (blue curves). Left plot shows loci 11kb from the centromere and the right shows loci 90 kb from the centromere. Only loci on chromosomes larger than 400 kb excluding Chr XII were included. The inset shows a 2D slice of the centromere positions for the loose (blue) and rigid (red) centromeres. Bonds are shown in grey and scale bar is 500 nm. **(C)** Effect of increasing polymer stiffness from one leading to a persistence length of 70 nm (red curves) to one with an angular spring constant that is 20% higher (blue curves). Loci are as in (B). Inset shows a 2D conformation of Chr I in the two conditions, scale bar is 500 nm. **(D)** MSD for loci 11 kb away from the centromere when comparing effects of increasing flexibility (blue and red curves, same as left panel of (B)) and combining flexibility and stiffness (grey curve).

We then modified variables related to microtubule tethering and chromatin stiffness and analyzed simulated chromatin dynamics. We first investigated the effect of weakening microtubule tethering to the kinetochore. The spring constant of the harmonic bond between the centromere and the SPB was loosened from a value ensuring a fixed bond to one yielding an energy of 1kbT at 105 nm extension from the 400 nm rest length. Loosening the attachment increased mobility for loci 11 kb away from the centromere, as we observed when microtubules were depolymerized by nocodazole, but left loci 90 kb unchanged **(Figure 2D**, **Figure 4B** - **Figure supplement 7**), indicating that tethering by microtubules induces only a local effect on chromatin mobility (see discussion). We then modified chromatin state by increasing global chromatin stiffness. In contrast to the position-dependent change when weakening microtubule tethering, this led to a global mobility increase independent of the distance from the centromere (**Figure 4C - Figure supplement 8**). This is expected, as even in the simple Rouse model, mobility increases when the spring constant between monomers increases (Herbert et al., 2017).

We reasoned that both effects could be at play in the pericentromeric region since the Smc5-2KE mutation displayed a greater increase in mobility than the nocodazole treatment. To test if this was possible, we combined a stiffness increase with the loosened chromatin tethering to microtubules. We observed that stiffening further increased chromatin dynamics, like the results seen *in vivo* for the difference between WT nocodazole and the *smc5-2KE* mutant (**Figure 2D**, **Figure 4D - Figure supplement 9**). These results show that a relaxation of microtubule attachment and a stiffer chromatin are sufficient to explain the increase in pericentromeric chromatin dynamics observed *in vivo* in the *smc5-2KE* mutant. They further support our hypothesis that Smc5 binds to both microtubules and chromatin to maintain pericentromeric chromatin folding and dynamics.

### The *smc5-2KE* mutant alters the mitotic spindle integrity

A correlation between pericentromeric chromatin compaction and spindle elongation has been reported previously, both in yeast and mammals (Bouck and Bloom, 2007; Mora-Bermúdez et al., 2007; Stephens et al., 2011). Since our results point to a role of the association between the Smc5/6 and microtubules on the dynamics and properties of pericentromeric chromatin, this prompted us to investigate spindle elongation. We first studied spindle length by measuring the distance between each spindle pole body (SPB) in dividing G2/M cells. We used SPB protein Spc42 fused to mCherry (He et al., 2000) to calculate the 2D distances between the two fluorescent centers of mass as above (**Figure 3**, **Figure 5A left**). In WT cells, spindle length average was 1.18 ± 0.29 µm, whereas *smc5-2KE* mutant exhibited a significant increase to 1.32 ± 0.33 µm (p value < 0,0001) (**Figure 5A left**), suggesting that the binding between Smc5 and microtubules contributes to the regulation of the spindle length.

**Figure 5:**
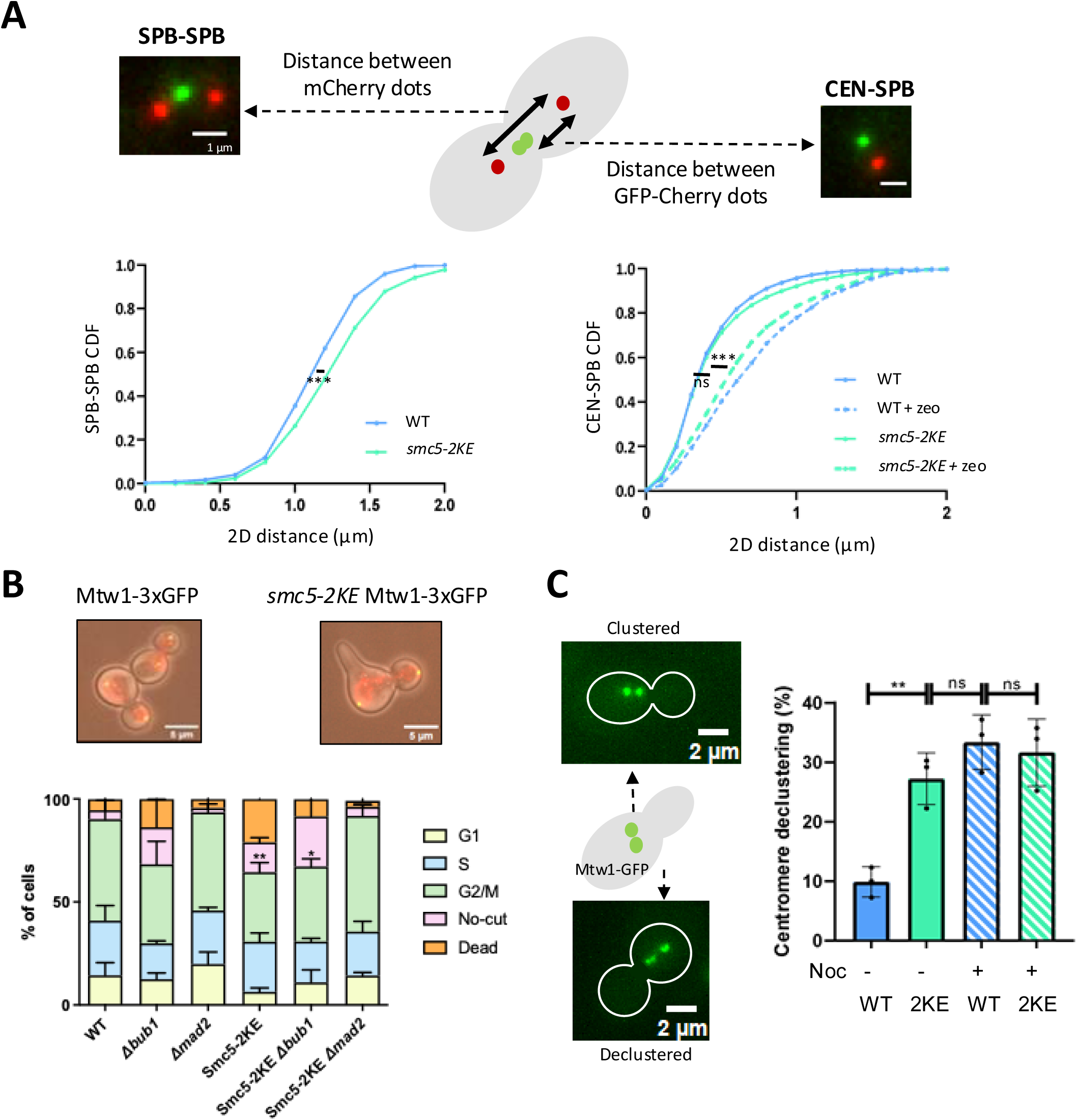
The *smc5-2KE* mutant alters the mitotic spindle integrity. **(A)** CEN IV was tagged with GFP and SPB with Spc42-mCherry. 2D distances between two SPBs (red dots) (**left**) and between Cen (green) and SPB (red) in G2/M cells (**right**) were measured in three independent biological replicates., Number of cells for SPB-SPB (N_WT_=224, N_2KE_=332) and for CEN-SPB (N_WT_=247, N_2KE_=380). As a positive control, CEN-SPB was measured in the presence of Zeocin with N_WT ZEO_=1299, N_2KE ZEO_=2033. Wilcoxon rank-sum test was used to analyze differences between groups, with “ns” for not significant and *** for P<0.0001. **(B)** Examples of cycling and defective mitotic cells are shown (top) and cycle phase distribution quantification (bottom) in *Mtw1-3xGFP* (WT) and *Smc5-2KE Mtw1-3xGFP* (2KE) cells, in the presence or not of deletion of *Mad2 (Δmad2*) or *Bub1* (*Δbub1*). Cells were categorized into G1, S, or G2/M phases. Additionally, dead cells and those exhibiting no-cut phenotype (no-cut) were counted in three experimental replicates N number of cells were counted N_WT_=439, N_WT NOC_=588, N_2KE_=413, N_2KE NOC_=652. T-test was used to analyze with * for P<0.01 and ** for P<0.001. **(C)** Mtw1 was labelled with 3xGFP to measure kinetochore clusters with or without nocodazole addition (15 ng/µl). Percentage of non-clustered cells are plotted. T- test was used to analyze differences between groups, with “ns” for not significant, ** for (P<0.001).

Alterations in spindle length can arise from changes in the length of the kinetochore microtubules connecting centromeres to the SPB, or modifications in the kinetochore/pericentromere itself. Measuring the distance between GFP-tagged CEN IV and Spc42-mCherry showed no difference between the WT and *smc5-2KE* mutant (**Figure 5A right**), suggesting that *smc5-2KE* does not alter the kinetochore microtubules length, while Zeocin control increased CEN-SPB distance as already demonstrated (**Figure 5A right**) (Hauer et al., 2017; Herbert et al., 2017). To then evaluate the kinetochore integrity in the *smc5-2KE* mutant, we measured centromere clustering *in vivo* by visualizing kinetochore protein Mtw1 fused to 3xGFP at the endogenous locus (Strecker et al., 2016). Mtw1 is a kinetochore-specific protein, part of the MIND complex, which links the inner and outer layers of the kinetochore (Maskell et al., 2010). While Mtw1-3xGFP expressing cells cycled normally, combination with *smc5-2KE* surprisingly led to a mitosis defect reminiscent of the no-cut phenotype, where completion of cytokinesis is delayed (**Figure 5B right**) (Norden et al., 2006). Strikingly, deletion of Mad2 or bub1, key spindle assembly checkpoint (*SAC*) components, suppressed the mitotic defect we observed, allowing cells to progress through mitosis despite the presence of the *Smc5-2KE* mutation and *Mtw1-3xGFP* (**Figure 5B right**).

In WT cells, Mtw1-3xGFP formed a single fluorescent focus due to the centromeric clustering typical of the Rabl configuration. Treatment with the microtubule-depolymerizing drug nocodazole induced fragmentation of the Mtw1-3xGFP labelling into several foci, consistent with centromeres declustering, as previously reported (Richmond et al., 2013; Strecker et al., 2016). Strikingly, when compared to WT, the *smc5-2KE* mutant presented a high percentage of cells with centromere declustering (27.26 ± 4.3% and 9.89 ± 2.5% for *smc5-2KE* and WT cells respectively, **Figure 5C**).

Overall, our results indicate that *smc5-2KE*, in the presence of Mtw1-3xGFP hypomorphic allele, exacerbates a kinetochore defect that activates the SAC. It suggests that the molecular link between chromatin and microtubules provided by Smc5/6 participates in kinetochore maintenance and mitotic spindle integrity.

### The *smc5-2KE* mutant affects homologous recombination at pericentromeres

To investigate whether Smc5/6 microtubule-dependent pericentromeric folding is required for genomic integrity, we induced targeted unique DSB in the pericentromere or far from it and analyzed its efficiency of repair by HR or NHEJ both in WT and mutated smc5-2KE strains according to (García Fernández et al., 2022a, 2021) (**Figure 6A**). I-*Sce*I is expressed on a plasmid with the galactose-inducible *GAL1-10* promoter to induce the DSB (+DSB), while an empty vector serves as control (-DSB). In the presence of a donor, successful HR leads to mCherry expression, detectable by red fluorescence in living cells (**Figure 6A**). Recipient sequence was introduced at 5 kb (named C) and 404 kb (named L) from CEN IV in both the WT and the *smc5-2KE* mutant strains, resulting in four distinct strains (**Figure 6B**). The I-*Sce*I cleavage efficiency in these strains, quantified by qPCR, was ∼80%, after 6 hours of I-*Sce*I induction, irrespective of the position of the cutting site, as previously observed (García Fernández et al., 2022a) (**Figure supplement 10A**). Cell survival was assessed by colony forming unit (CFU) assays. As expected, in the absence of I-*Sce*I, WT and *smc5-2KE* C and L colonies grew in both inducing and non-inducing conditions (**Figure 6B**).

**Figure 6:**
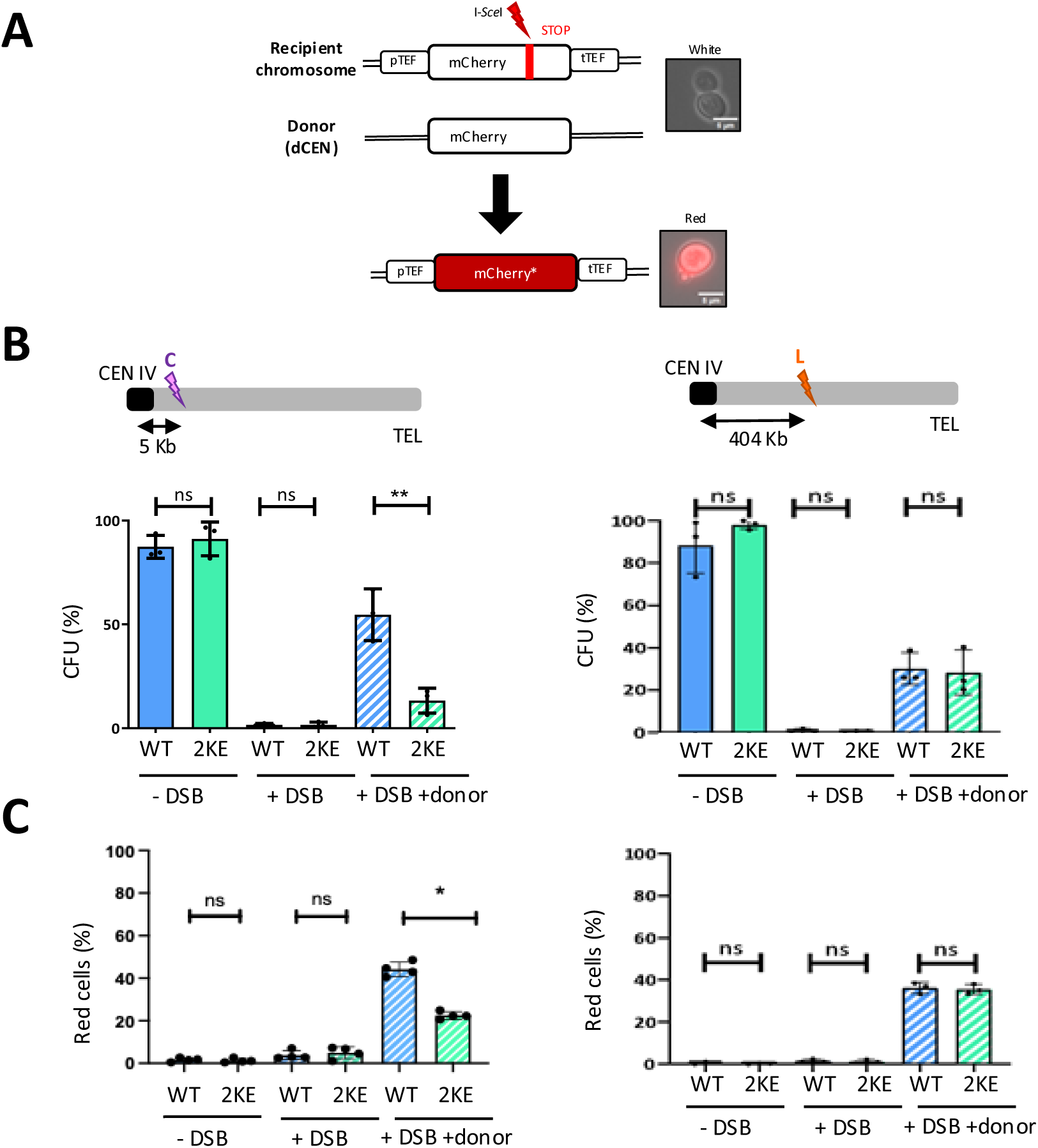
The *smc5-2KE* mutant decreases homologous recombination efficiency at pericentromeres. **(A)** Schematic representation of the system to track homologous recombination (HR). In the recipient mCherry, I-*Sce*I has a 30 bp I-*Sce*I cutting site sequence followed by a stop codon (red bar) inserted at 57 amino acids from the end of the mCherry coding sequence. The donor (dCen) is a full-length mCherry lacking promoter and terminator sequences, carried on a centromeric plasmid. A functional mCherry could be expressed only after successful HR. **(B)** Colony formation unit (CFU) test showing three conditions: without DSB (-DSB), with a DSB (+DSB) and with a DSB and a donor (DSB+donor) in the C and in the L strain for the WT and for the *smc5-2KE* mutant. Ratio between Induced I-*Sce*I/Non induced I-*Sce*I. Three experiments were conducted with the number of cells counted as follows: For the C strain, N_WT PRS_ =487, N_WT PB_ =528, N_WT Cen_ =668, N_2KE PRS_ =330, N_2KE PB_ =620, N_2KE Cen_ =752. For the L strain, N_WT PRS_ =344, N_WT PB_ =309, N_WT Cen_ =328, N_2KE PRS_ = 261, N_2KE PB_ =379, N_2KE Cen_ =387. Mann-Whitney test was used to analyse differences between groups, with “ns” for not significant (P>0.05) and ** for P<0.001. **(C)** Number of red cells (%) without DSB (-DSB), with a DSB (+DSB) and with a DSB and a donor (DSB+donor) in the C and in the L strain. Three experiments were conducted, counting the number of cells for the C strain as follows: N_WT PRS_ =707, N_WT PB_ =719, N_WT Cen_ =2180, N_2KE PRS_ = 727, N_2KE PB_ =660, N_2KE Cen_ =2553. For the L strain, the counts were N_WT PRS_ =506, N_WT PB_ =607, N_WT Cen_ =1299, N_2KE PRS_ =744, N_2KE PB_ =712, N_2KE Cen_ =1551. Mann-Whitney test was used to analyse differences between groups, with “ns” for not significant (P>0.05) and * for P<0.005.

Interestingly, providing the donor sequence on the dCEN plasmid to allow DSB repair by HR, had different outcomes in *smc5-2KE* depending on the DSB position. When the DSB was in the pericentromeric region, the *smc5-2KE* mutant showed significantly reduced survival rates compared to the WT (C strains, 15 ± 6% and 55 ± 12.4%, respectively, **Figure 6B**). In contrast, both WT and the *smc5-2KE* mutant presented similar survival rates for a DSB located in the L position (40 ± 7.5% for both L strains) (**Figure 6B**). The smc5-2KE mutant generated fewer mCherry-positive cells than the WT at the pericentromeric DSB (WT, 44 ± 3.4% and *smc5-2KE* 22.5 ± 1.7%. Importantly, at the L position, both WT and smc5-2KE strains showed comparable mCherry expression levels (36 ± 2.5% for both). HR defect could not be explained by a loss of the centromeric plasmid (**Figure supplement 10B**) nor a change in Smc5/6 occupancy at DSB, as ChIP-qPCR of Smc5-MYC or *smc5-2KE*-MYC revealed no significant differences, irrespective of the DSB position (**Figure supplement 10C-D**).

Overall, these results indicate that *smc5-2KE* impairs HR at pericentromeres, but not along the chromosome arm, and suggest that pericentromeric folding due to Smc5 binding to microtubules is required for pericentromeric integrity.

Local mobility induced by a DSB is facilitated by microtubules (Lawrimore et al., 2017; Oshidari et al ., 2018) and has been proposed to promote DSB repair by HR (Miné-Hattab et al., 2017; Neumann et al., 2012). We therefore analyzed chromatin mobility at the DSB in the *smc5*-*2KE* mutant. We introduced a lacO-LacI-GFP array at 10 kb away from the I-*Sce*I cutting site in WT and mutant C and L strains and verified by qPCR that the efficiency of I-*Sce*I cutting was not affected by the array in all strains (80% at 6 hours after induction) (**Figure supplement 10E**). Chromatin dynamics was tracked after 6 hours of induction, and MSD was plotted through increasing time intervals (**Figure supplement 10F-G**). DSB induction in WT led to an increase in chromatin dynamics, independently of the DSB position, whereas in the *smc5*-*2KE* mutant, DNA damage did not cause an additional increase in mobility on top of the one generated by the mutation, as reported above (**Figure 1** -**figure supplement 10F-G**). These results show that differences in HR efficiency between a pericentromeric DSB and a mid-arm DSB are not related to changes in local chromatin dynamics and suggest that Smc5-microtubule association is required for a fraction of the mobility induced by a DSB.

## Discussion

Pericentromeres across eukaryotic evolution are enriched with SMC complexes, which share the conserved role of extruding DNA to create loops (Davidson et al., 2019; Ganji et al., 2018; Pradhan et al., 2023). The pericentromeric enrichment of SMCs and their extruding activity theoretically leads to a structure described, from a physical perspective, as a bottlebrush, balancing the outward pulling forces of microtubule with the inward attractive forces of chromatin during cell division (Bloom, 2024). Smc5/6 has been shown to associate with microtubules in budding yeast and *Xenopus* (Gache et al., 2010; Gutierrez-Escribano et al., 2020; Laflamme et al., 2014 and our present data - **Figure supplement 1**). However, the role of this association in organizing pericentromeric structure *in vivo* was previously unknown. By measuring chromatin dynamics in the *smc5-2KE* mutant, which decreases the affinity between Smc5 and microtubules, we demonstrate that Smc5 associates with microtubules *in vivo* and regulates chromatin mobility along the chromosome, as well as chromatin properties in the pericentromeric region. Moreover, we show that Smc5/6 association with microtubules specifically maintains the integrity of the mitotic spindle and contributes to DSB repair efficiency in the pericentromeric region.

We provide evidence that Smc5/6 offers a molecular link to microtubules through the conserved residues K624 and K631 in the hinge region of Smc5. First, alphafold3 modelling shows a high probability of interaction between extruded Smc5 hinge and alpha/beta dimers of tubulin (**Fig 1A**). Second, the microtubule binding co-sedimentation assay with Smc5 is decreased when K624 and K631 are mutated to aspartic acid (**Figure 1C**). Third, in the *smc5-2KE* mutant, chromatin dynamics mirrors microtubule depolymerization (**Figure 2D,F**) and resembles the effects observed in the *ndc80-1* kinetochore mutant (**Figure 2E,F**). The ability of Smc5/6 to bind microtubules is thus an important emerging property of this complex.

The association between Smc5/6 and microtubules restricts chromatin mobility and has critical consequences for the folding of the pericentromeric region. In this region, measurements of both chromatin mobility and intra-chromosomal distances in the *smc5-2KE* mutant indicate that reduced association with microtubules, and only them, impacts chromatin structure, potentially leading to increased rigidity or decompaction, as shown by polymer modelling (**Figure 4D**). Notably, changes in nuclear environment (i.e. changes in chromatin density because of centromere release in a configuration where centromeres are clustered due to Rabl configuration) are consistent with an increased chromatin mobility with similar alpha exponent of the MSD **(Fig 1E).** How could the Smc5-2KE mutation affect the properties of pericentromeric chromatin? Loss of microtubules polymerization or kinetochore mutants, such as Gal-Nuf2, where Nuf2 is repressed in the presence of glucose, result in decreased levels of histone H4 and H2B in the pericentromere, as shown by FRAP (Verdaasdonk et al., 2012). Because histone reduction has been linked to chromatin decompaction and increased chromatin dynamics (Amitai et al., 2017; Cheblal et al., 2020; Hauer et al., 2017), disruption of microtubule-induced tension in the *smc5-2KE* mutant could similarly result in decreased histones occupancy, chromatin decompaction and subsequent chromatin mobility increase. The fact that the loss of microtubules also increases intra-chromosomal contacts and affects pericentromeric configuration (Paldi et al., 2020), suggest that the tension created by microtubules can regulate chromatin structure close to the attachment sites (Paldi et al., 2020). Accordingly, the interaction between Smc5 and microtubules, disrupted in the *smc5-2KE*, may contribute to microtubule-induced tension and spindle robustness, thereby influencing pericentromeric chromatin architecture.

Another interesting possibility by which Smc5/6 might regulate pericentromeric chromatin structure could be via its DNA compaction activity, as recently shown by single molecule experiments with purified human and yeast Smc5/6 (Gutierrez-Escribano et al., 2020; Serrano et al., 2020). Mechanistically, the altered chromatin structure in the *smc5-2KE* mutant could result from impaired loop extrusion by Smc5/6 dimers (Pradhan et al., 2023). However, we show here that acute depletion of the entire complex - thereby abolishing loop extrusion activity - has an impact on chromatin mobility opposite to the Smc5-2KE mutation (**Figure supplement 6)**. This finding strongly suggests that loop extrusion is not responsible for the altered chromatin behavior in *smc5-2KE.* Instead, Smc5/6 could promote chromatin compaction through alternative mechanisms, such as microtubule-associated nucleosome stabilization. Defects in cohesin or condensin recruitment could also contribute to pericentromeric decompactionr or stiffness leading to increased chromatin dynamics (**Figure 1C**) (Cheblal et al., 2020; Dion et al., 2013; Verdaasdonk et al., 2013). Pairing defects by cohesin appear unlikely in the *smc5-2KE* mutant, as cohesion is intact, contrasting what is observed upon cohesin-depleted cells (**Figure supplement 6D,E).** Overall, our observations suggest a pathway in which microtubule forces transmitted via Smc5/6 regulate pericentromeric chromatin folding and stiffness, independently of loop extrusion or cohesion.

It is intriguing that disrupting the microtubule binding to centromeres, either by depolymerization of microtubules in the presence of nocodazole, by affecting the kinetochore in the *ndc80-1* mutant or with the *smc5-2KE* mutant results in increased chromatin dynamics not only close to the centromere but also, to a lesser extent, at locations distant from it. It is possible that once centromeres are freed from microtubules, they exhibit increased movement, which could be translated into greater chromatin mobility further along the chromosome. By analogy, a rope with movement at one end could lead to increased movement at the other end. However, by simulating weakened microtubule anchoring, the polymer model fails to fully reproduce the *in vivo* increase in chromatin dynamics away from the centromere (**Figure 4B**), possibly due to unmodeled features such as SMC-mediated loop formation or centromeres declustering, which might influence chromatin density *in vivo*.

One of the pericentromeric chromatin functions mediated by the association of the Smc5/6 complex with microtubules is clarified by the "no-cut" phenotype (**Figure 5B**). Defects in cytokinesis are indeed observed when the *smc5-2KE* mutant is combined with Mtw1-3xGFP. This phenotype, also described in cells with mutations in kinetochore proteins such as *ndc10-1* and *ndc80-1* alleles (Bouck and Bloom, 2005; Norden et al., 2006) suggests that the interaction between the Smc5-2KE mutation and Mtw1-3xGFP is associated with compromised kinetochores (Fujioka et al., 2002). The “no-cut” phenotype has been described in cells with incomplete DNA segregation (Mendoza et al., 2009), indicating that disruption in chromosome segregation can lead to cytokinesis defects. Strikingly, deletion of *Mad2* or *Bub1* suppressed the mitotic defect, indicating that the no-cut phenotype is SAC-dependent and further supports the hypothesis that Smc5-2KE and Mtw1-GFP exacerbate kinetochore dysfunction to a degree that activates the SAC. Thus, it is conceivable that the combination of Smc5-2KE with Mtw1-3xGFP results in chromosome segregation defects leading to the “no-cut” phenotype and pinpoints to the important function of Smc5/6 in establishing pericentromeric chromatin through its association with microtubules.

It has previously been shown both in budding yeast and mammalian cells, that Smc5/6 is enriched around DSBs (De Piccoli et al., 2006; Lindroos et al., 2006; Potts et al., 2006) and contributes to HR by resolving Holliday junctions (Bermúdez-López et al., 2010; Pebernard et al., 2006; Torres-Rosell et al., 2005), as well as by maintaining the ends of a DSB together (Phipps et al., 2024). In line with the role of Smc5/6 in HR, we show that the Smc5-2KE mutation leads to reduced repair efficiency only at pericentromeres (**Figure 6**). HR efficiency could be impacted either because pericentromeric chromatin properties are modified as we observed due to the lack of association between Smc5/6 and microtubules (**Figure 3**), or because *smc5-2KE* mutant directly impairs pericentromeric DSB repair. For instance, mutating the hinge domain of Smc5, which has been shown to have affinity for ssDNA *in vitro* (Alt et al., 2017; Roy et al., 2011), could affect the putative interaction between the Smc5/6 complex and ssDNA at DSB, potentially impacting 5’ DNA resection at DSBs or the maintenance of the DSB ends (Phipps et al., 2024). Alternatively, the lack of association between microtubules and the Smc5/6 could hinder the efficient relocalization of damaged pericentromeres to the nuclear periphery, in line with the reported role of the Smc5/6 complex relocalization for DSBs in repetitive rDNA sequences in budding yeast (Torres-Rosell et al., 2007) and in heterochromatin in *Drosophila* (Caridi et al., 2018; Ryu et al., 2015). Furthermore, upon a DSB near the centromere, DNA resection may lead to detachment of the centromere from microtubules, triggering the formation of damage-induced microtubules (DIM) to recapture the centromere as proposed (Oshidari et al., 2018). This process would require Smc5/6; thus, a mutation affecting association with DIMs could hamper centromere recapture and efficient DSB repair. These alternatives explanation deserve further investigation in future studies.

Taken together, our focus on chromatin dynamics emerged as a powerful approach for understanding the functional relevance of the association between Smc5/6 with microtubules on the maintenance and integrity of pericentromeric chromatin. We propose that the enrichment of Smc5/6 at pericentromeres, together with its association with microtubules, contribute to the maintenance of pericentromere chromatin stability in the face of mitotic forces. Smc5/6 enrichment in the pericentromeric region is conserved throughout mitosis of many organisms (Jeppsson et al., 2014; Lindroos et al., 2006; Venegas et al., 2020). It is likely that a similar association between Smc5/6 and microtubules occurs in mammals during open mitosis, the only phase when microtubules interact with centromeric regions in mammals. Therefore, the role of Smc5/6 in preserving pericentromeric structure during mitosis might be conserved through evolution.

## Material and methods

### Yeast strains

*S. cerevisiae* strains used in this study are detailed in **Table 1** and isogenic to BY4741.

**Table 1:** genotypes of used strains.

| Common name | Strain | Genotype | Reference |
| --- | --- | --- | --- |
| **11kb Chr IV** | YPE4019 | Mata, ura3, trp1-1, leu2-3, his3-11,15, can1-100, ade2-1, P<sub>MET3</sub>-CDC20::URA3, Spc42-tdTomato::Nat, leu2:: P<sub>URA3</sub>-TetR-GFP::LEU2, tetOx224-URA3 (tetOs 11,5kb to right of CEN4), yil015w::KanMX | Paldi *et al* 2020 |
| **11kb-2KE Chr IV** | YPE4144 | Mata, ura3, trp1-1, leu2-3, his3-11,15, can1-100, ade2-1, P<sub>MET3</sub>-CDC20::URA3, Spc42-tdTomato::Nat, leu2:: P<sub>URA3</sub>-TetR -GFP::LEU2, tetOx224-URA3 (tetOs 11,5kb to right of CEN4), yil015w::KanMX, YOL034W::K624E, YOL034W::K631E | This study |
| **23kb Chr IV** | YEF1659 | Mata, ura3, trp1-1, leu2-3, his3-11,15, can1-100, ade2-1, P<sub>MET3</sub>-CDC20::URA3, Spc42-tdTomato::Nat, leu2:: P<sub>URA3</sub>-TetR -GFP::LEU2, tetOX224-URA3 (tetOs 11,5kb to right of CEN4) | Paldi *et al* 2020 |
| **23kb-2KE Chr IV** | YPE4145 | Mata, ura3, trp1-1, leu2-3, his3-11,15, can1-100, ade2-1, P<sub>MET3</sub>-CDC20, Spc42-tdTomato::Nat, leu2:: P<sub>URA3</sub>-TetR -GFP::LEU2, tetOX224-URA3 (tetOs 23kb to right of CEN4), YOL034W::K624E, YOL034W::K631E | This study |
| **90kb Chr IV** | YEF1081 | Mat alpha, ura3Δ0 or-851, his3Δ1, can1::HPHMX, his3::HIS3-LacI-GFP, YDR042C::lacO-LEU2 | This study |
| **90kb-2KE Chr IV** | YEF1654 | Mat alpha, ura3Δ0 or-851, his3Δ1, can1::HPHMX, his3::HIS3-LacI-GFP YDR042C::lacO-LEU2, YOL034W::K624E, YOL034W::K631E | This study |
| **730kb Chr IV** | YEF1068 | Mat alpha, ura3Δ0, leu2Δ0, his3Δ1, lys2Δ202, ade2-661, iyGL117::TetR-mRFP-NATMX, YDR354W::tetO-URA3 HIS3::HIS3-LacI-GFP YDR259C::lacO-TLEU2 | This study |
| **730kb-2KE Chr IV** | YEF1706 | Mat alpha, ura3Δ0, leu2Δ0, his3Δ1lys2Δ202, ade2-661, iyGL117::tetR-mRFP NATMX YDR354W::tetO-URA3 HIS3::HIS3-LacI-GFP, YDR259C::lacO- LEU2, YOL034W::K624E, YOL034W::K631E | This study |
| **90kb-2KR Chr IV** | YPE4180 | Mat alpha, ura3Δ0, trp1Δ63, leu2Δ0, his3Δ1, lys2Δ202, can1::HPHMX, ade2-661, lys2::TetR-GFP-LYS2, yel053c ::KanMX::tetO-URA3, Chr IV 453880+453959 ::KanMX::mCherry*-IScl(cs), YOL034W::K624R, YOL034W::K631ER | This study |
| **90kb Chr V** | YEF1454 | Mat alpha, ura3Δ0, trp1Δ63, leu2Δ0, his3Δ1, lys2Δ202, can1::HPHMX, ade2-661, lys2::TetR-GFP-LYS2, yel053c ::KanMX:: tetO-URA3, Chr IV 453880+453959 ::KanMX::mCherry*-IScl(cs) | García-Fernandez *et al.*, 2022 |
| **90kb-2KE Chr V** | YEF1666 | Mat alpha, ura3Δ0, trp1Δ63, leu2Δ0, his3Δ1, lys2Δ202, can1::HPHMX, ade2-661,::TetR-GFP-LYS2, yel053c ::KanMX:: tetO-URA3, Chr IV 453880+453959 :: KanMX::mCherry*-IScl(cs), YOL034W::K624E, YOL034W::K631E | This study |
| **SMC6-AID AFB2** | YPE4059 | Mat a, trp1-1, leu2-3, his3-11, ade2-1, P<sub>MET3</sub>-CDC20::URA3, Spc42-tdTomato::Nat, leu2::PURA3::tetR-GFP::LEU2, tetOX224-URA3 (tetOs 11,5kb to right of CEN4), Smc5-AID-9myc-KanMX, his3-11,15 :: pEF604 (=AFB2-HIS3) | This study |
| **SMC1-AID AFB2** | YPE4173 | Mat a, trp1-1, leu2-3, his3-11, ade2-1, P<sub>MET3</sub>-CDC20::URA3, Spc42-tdTomato::Nat, leu2::PURA3::tetR-GFP::LEU2, tetOX224-URA3 (tetOs 11,5kb to right of CEN4), yil015w::KanMX, his3-11,15::AFB2-HIS3(pEF604), Smc6-AID-9myc-TRP1 | This study |
| **SMC5-AID AFB2** | YPE4176 | Mat a, trp1-1, leu2-3, his3-11, ade2-1, P<sub>MET3</sub>-CDC20::URA3, Spc42-tdTomato::Nat, leu2::PURA3::tetR-GFP::LEU2, tetOX224-URA3 (tetOs 11,5kb to right of CEN4), yil015w::KanMX, his3-11,15::AFB2-HIS3(pEF604), Smc1-AID-3myc-TRP1 | This study |
| **Rad52 GFP** | YEF651 | Mata, ura3Δ0, leu2Δ0, his3Δ1, met15Δ0, YML032C-GFP-HIS3MX6 | |
| **Rad52-GFP 2KE** | YEF1647 | Mata, ura3Δ0, leu2Δ0, his3Δ1, met15Δ0, YML032C-GFP-HIS3MX6, YOL034W::K624E, YOL034W::K631E | This study |
| **RAD53-HA** | YEF954 | Mat alpha, ura3-Δ851, trp1-Δ63, leu2Δ1, his3Δ200, ade2-661, yml032c::KANMX, rad53-HA | |
| **RAD53-HA 2KE** | YEF1882 | Mat alpha, ura3-Δ851, trp1-Δ63, leu2Δ1, his3Δ200, ade2-661, yml032c::KANMX, rad53-HA, YOL034W::K624E, YOL034W::K631E | This study |
| **R1** | YEF1065 | Mat a, ura3Δ0, leu2Δ0/, his3Δ1, lys2Δ202, ade2-661, can1::HPHMX, iyGL117::tetR-mRFP- NATMX, YDR003W::tetO-TEF-URA, HIS3::HIS3-LacI-GFP, YDR095C::256lacO-TEF-LEU2 | Herbert *et al* 2017 |
| **R1 2KE** | YEF1887 | Mat a, ura3Δ0, leu2Δ0/, his3Δ1, lys2Δ202, ade2-661, can1::HPHMX, iyGL117::tetR-mRFP- NATMX, YDR003W::tetO-TEF-URA, HIS3::HIS3-LacI-GFP, yDR095c::256lacO-LEU2, YOL034W::K624E, YOL034W::K631E | This study |
| **R3** | YEF1068 | Mat alpha, ura3Δ0, leu2Δ0, his3Δ1, lys2Δ202, ade2-661, iyGL117::tetR-mRFP-NATMX, YDR354W::tetO-TEF-URA3 HIS3::HIS3-LacI-GFP YDR259C::lacO- LEU2 | Herbert *et al* 2017 |
| **R3 2KE** | YEF1706 | Mat alpha, ura3Δ0, leu2Δ0, his3Δ1, lys2Δ202, ade2-661, iyGL117::tetR-mRFP NATMX YDR354W::tetO-TEF-URA3 HIS3::HIS3-LacI-GFP YDR259C::lacO- LEU2, YOL034W::K624E, YOL034W::K631E | This study |
| **Tub2-GFP** | YPE257 | Mata, ura3-52, trp1Δ63, leu2Δ1, lys2-801, ade2-661, ura3-52::TUB2-GFP-URA3 | |
| **Tub2-GFP SMC5-2KE** | YEF1864 | Mata, ura3-52, trp1Δ63, leu2Δ1, lys2-801, ade2-661, ura3-52::TUB2-GFP-URA3, YOL034W::K624E, YOL034W::K631E | This study |
| **Smc5-ndc80-1** | YEF1928 | Mat alpha, ura3Δ0, trp1Δ63, leu2Δ0, his3Δ1, lys2Δ202, can1::HphMX,ade2-661, lys2::TetR-GFP-LYS2, yel053c ::KanMX:: tetO-URA3, Chr IV 453880+453959::KanMX:: mCherry*-I-SceI(cs), ndc80-1 | This study |
| **ndc80-1 Smc5-2KE** | YEF1930 | Mat alpha, ura3Δ0, trp1Δ63, leu2Δ0, his3Δ1, lys2Δ202, can1::HphMX,ade2-661/,lys2::TetR-GFP-LYS2, yel053c ::KanMX:: tetO-URA3, Chr IV 453880+453959::KanMX:: mCherry*-I-SceI(cs), YOL034W::K624E, YOL034W::K631E, ndc80-1 | This study |
| **L WT sDSB** | YEF1203 | Mat alpha, ura3Δ0, trp1Δ63, leu2Δ0, his3Δ1, lys2Δ202, ade2-661, lys2::TetR-GFP-LYS2, can1::HphMX, YEL053C :: KANMX:: tetO-URA Chr IV 853996+854075:: KANMX:: mCherry*-I-SceI(cs) | García-Fernandez *et al.*, 2022 |
| **L 2KE sDSB** | YPE4131 | Mat alpha, ura3Δ0, trp1Δ63, leu2Δ0, his3Δ1, lys2Δ202, ade2-661, lys2::TetR-GFP-LYS2, can1::HPHMX, YEL053C::KanMX::tetO-URA, Chr IV 853996+854075:: KanMX:: mCherry*-I-SceI(cs), YOL034W::K624E, YOL034W::K631E, | This study |
| **C - DSB Local mobility** | YPE4132 | Mata,ura3Δ0,,leu2Δ0/,his3Δ1/, met15Δ0/,YOL034W-13xMYC-URA3, Chr IV 453880+453959::KanMX::mCherry*-I-SceI(cs), ymr008C::KanMX::lacO-LEU2, LacI-HIS3, HIS3-STOP, p2u-HIS3 | This study |
| **C + DSB Local mobility** | YPE4133 | Mata, ura3Δ0, leu2Δ0, his3Δ1, met15Δ0, YOL034W-13xMYC-URA3, Chr IV 453880+453959::KANMX::mCherry*-I-SceI(cs), ymr008C::kanMX::lacO-LEU2, LacI-HIS3, HIS3-STOP, pGALISceI-HIS3 | This study |
| **C 2KE - DSB Local mobility** | YPE4136 | Mata, ura3Δ0, leu2Δ0, his3Δ1, met15Δ0, YOL034W::K624E, YOL034W::K631E, YOL034W-13xMYC-URA3, Chr IV 453880+453959::KanMX: mCherry*-I-SceI(cs), ymr008C::kanMX::lacO-LEU2, LacI-HIS3, HIS3-STOP, p2u-HIS3 | This study |
| **C 2KE + DSB**<br>**Local**<br>**mobility** | YPE4137 | Mata, ura3Δ0, leu2Δ0, his3Δ1, met15Δ0, YOL034W::K624E, YOL034W::K631E, YOL034W-13xMYC-URA3, Chr IV 453880+453959::KanMX::mCherry*-I-SceI(cs), ymr008C::kanMX::lacO-LEU2, LacI-HIS3, HIS3-STOP, pGALISceI-HIS3 | This study |
| **L - DSB sDSB**<br>**Local**<br>**mobility** | YPE4134 | Mata, ura3Δ0, leu2Δ0, his3Δ1, met15Δ0, YOL034W-13xMYC-URA3, Chr IV 853996+854075::KanMX::mCherry*-I-SceI(cs), ydr193w::kanMX::lacO-LEU2, LacI-HIS3, HIS3-STOP, p2u-HIS3 | This study |
| **L + DSB Local**<br>**mobility** | YPE4135 | Mata, ura3Δ0, leu2Δ0, his3Δ1, met15Δ0, YOL034W-13xMYC-URA3, Chr IV 853996+854075::KanMX::mCherry*-I-SceI(cs), ydr193w::kanMX::lacO-LEU2, LacI-HIS3, HIS3-STOP, pGALISceI-HIS3 | This study |
| **L 2KE - DSB**<br>**Local**<br>**mobility** | YPE4138 | Mata, ura3Δ0, leu2Δ0, his3Δ1, met15Δ0, YOL034W::K624E, YOL034W::K631E, YOL034W-13xMYC-URA3, Chr IV 853996+854075::KanMX::mCherry*-I-SceI(cs), ydr193w::kanMX::lacO-LEU2, LacI-HIS3, HIS3-STOP, p2u-HIS3 | This study |
| **L 2KE + DSB**<br>**Local**<br>**mobility** | YPE4139 | Mata, ura3Δ0, leu2Δ0, his3Δ1, met15Δ0, YOL034W::K624E, YOL034W::K631E, YOL034W-13xMYC-URA3, Chr IV 853996+854075::KanMX::mCherry*-I-SceI(cs), ydr193w::kanMX::lacO-LEU2, LacI-HIS3, HIS3-STOP, pGALISceI-HIS3 | This study |
| **C - DSB ChIP-**<br>**qPCR** | YEF1922 | Mata, ura3Δ0, leu2Δ0, his3Δ1, met15Δ0, YOL034W-13xMYC-URA3, Chr IV 853996+854075:: kanMX::mCherry*-ISceI(cs), p2u-HIS3 | This study |
| **C + DSB**<br>**ChIP-qPCR** | YEF1923 | Mata, ura3Δ0, leu2Δ0, his3Δ1, met15Δ0, YOL034W-13xMYC-URA3, Chr IV 853996+854075::kanMX::mCherry*-I-SceI(cs), pGALISceI-HIS3 | This study |
| **C 2KE - DSB**<br>**ChIP-qPCR** | YEF1934 | Mata, ura3Δ0, leu2Δ0, his3Δ1, met15Δ0, YOL034W::K624E, YOL034W::K631E, YOL034W-13xMYC-URA3, Chr IV 453880+453959 :: kanMX :: mCherry*-I-SceI(cs), p2u-HIS3 | This study |
| **C 2KE + DSB ChIP-qPCR** | YEF1925 | Mata, *ura3Δ0*, *leu2Δ0*, *his3Δ1*, *met15Δ0*, YOL034W::K624E, YOL034W::K631E, YOL034W-13xMYC-URA3, Chr IV 453880+453959 :: kanMX :: mCherry*-I-SceI(cs), P<sub>GAL1-10</sub>-I-SceI-HIS3 | This study |
| **L - DSB ChIP-qPCR** | YEF1898 | Mata, *ura3Δ0*, *leu2Δ0*, *his3Δ1*, *met15Δ0*, YOL034W-13xMYC-URA3, Chr IV 853996+854075:: kanMX :: mCherry*-I-SceI(cs), p2u-HIS3 | This study |
| **L + DSB ChIP-qPCR** | YEF1900 | Mata, *ura3Δ0*, *leu2Δ0*, *his3Δ1*, *met15Δ0*, YOL034W-13xMYC-URA3, Chr IV 853996+854075::kanMX::mCherry*-I-SceI(cs), P<sub>GAL1-10</sub>-I-SceI-HIS3 | This study |
| **L 2KE - DSB ChIP-qPCR** | YEF1926 | Mata, *ura3Δ0*, *leu2Δ0*, *his3Δ1*, *met15Δ0*, YOL034W::K624E, YOL034W::K631E, YOL034W-13xMYC-URA3, Chr IV 853996+854075::kanMX::mCherry*-I-SceI(cs), p2u-HIS3 | This study |
| **L 2KE + DSB ChIP-qPCR** | YEF1920 | Mata, *ura3Δ0*, *leu2Δ0*, *his3Δ1*, *met15Δ0*, YOL034W::K624E, YOL034W::K631E, YOL034W-13xMYC-URA3, Chr IV 853996+854075::kanMX::mCherry*-I-SceI(cs), P<sub>GAL1-10</sub>-I-SceI-HIS3 | This study |
| **CEN-SPB** | YEF1019 | Mat a, *his3Δ1* or *his3Δ200*, *lys2Δ202* or 0, cenIV-tetO-URA3, *lys2Δ202*::tetR-GFP-LYS2, SPC42-mCherry-HIS3 | García-Fernandez *et al.*, 2021 |
| **CEN-SPB Smc5-2KE** | YEF1691 | Mat a, *his3Δ1* or *his3Δ200*, *lys2Δ202* or 0, cenIV-tetO-URA3, *lys2Δ202*::tetR-GFP-LYS2, SPC42-mCherry-HIS3, YOL034W::K624E, YOL034W::K631E | This study |
| **Mtw1-3xGFP** | YEF1017 | Mat a, *ura3-1*, *trp1-1*, *leu2Δ3,112*, *his3-11*, *can1-100*, *ade2-1*, *ura3::TUB1-CFP::URA3*, YAL034W-A-3xGFP::HIS3, *yil015w::KANMX* | García-Fernandez *et al.*, 2021 |
| **Mtw1-3xGFP Smc5-2KE** | YEF4025 | Mat a, *ura3-1*, *trp1-1*, *leu2Δ3,112*, *his3-11*, *can1-100*, *ade2-1*, *ura3::TUB1-CFP::URA3*, YAL034W-A-3xGFP::HIS3, YOL034W::K624E, YOL034W::K631E, *yil015w::KANMX* | This study |
| **SMC5-3xSTII, CBP-9xHIS-SMC6** | D10079 | Mata, W303, ura3-1, trp1-1, leu2-3,112, his3-11,15, can1-100, ade2-1, [PSI+], rad5-535, SMC5-3xSTII::TRP1, CBP-9xHIS-SMC6 | This study |

### Strain construction

Mutations of K624E and K631E of *SMC5* were performed with CRISPR/Cas9 using a 80 pb donor carrying the desired mutations as described in (García Fernández et al., 2022a). Unique duplexed 20 nucleotides long primers next to the PAM sequence (AGG) close to K631 (oACS13/14) were cloned into a replicative Cas9-KANMX plasmid (pEF562, Addgene plasmid #100955) to yield (pEF590) verified by PCR and sequencing. Equimolar amounts of pEF590 and the 80 pb donor (oACS69) were used to transform the desired strains and the corresponding smc5-2KE transformants were verified by sequencing. The same strategy was used to obtain smc5-2KA and smc5-2KR with different donor sequences carrying the specific mutation (oACS97 and oACS270 respectively).

The strains used to track the local mobility of a DSB were constructed as described in (García Fernández et al., 2022a). Briefly, the I-*Sce*I cutting site was inserted at the desired position, by replacing a KANMX cassette integrated into intergenic regions of chromosome IV at positions 453880-453959 (C strain) or 853996-854075 (L strain) with an mCherry sequence containing the I-*Sce*I cutting site. LacO repeats at 10 kb from the cutting site were introduced by replacing a KANMX cassette integrated at positions 462252-462602 (strain C) or 844554-844952 (strain L) with pEF222 plasmid digested by *Bgl*II (BioLabs, R0144). LacI-GFP was introduced by digesting plasmid pEF569 (Therizols et al., 2010) with *Nhe*I (BioLabs, R3131). The ARS-CEN-HIS3 plasmid (pB07, obtained from B. Pardo) for I-*Sce*I expression under inducible pGAL1-10 promoter, was used after insertion of a stop codon in the *HIS3* genomic sequence by CrispR/Cas9 as above. The 20 nucleotides duplex oligos (oACS204/205) next to the PAM (TGG) sequence in *HIS3* and an 80 pb primer (oACS208) donor were used.

Depletion of *SMC5* was obtained by using a degron AFB2 based inducible (Nishimura and Kanemaki, 2014). AID-9MYC tag at the C-terminus of *SMC5* was introduced by homologous recombination using pEF565 (Morawska and Ulrich, 2013) and (oACS40/53) oligonucleotides.

AFB2 was introduced at the *HIS3* locus by digesting the plasmid pEF604 (obtained from T.Teixeira) carrying *AFB2-HIS3* by *Nhe*I (BioLabs, R3131).

### Media and growth conditions

Cells were grown in SC medium (2 g/l amino acids mix, 20 g/l sugar, 6.7 g/l yeast nitrogen base) until they reached the mid-log phase. Multiple DNA damages were induced with 250 µg/ml Zeocin (Thermo Fisher, R25001) added for 4 hours (Herbert et al., 2017). Microtubules were depolymerized using 15 ng/µl of nocodazole (Sigma, M1404) dissolved in DMSO, for 30 minutes. Depolymerization was checked in parallel by Tub1-GFP disappearance in strain YPT257 (Therizols et al., 2010). To induce a single DSB, 2% of galactose was added for 6 hours.

G1 arrest was performed using 50 ng/ml of alpha-factor (Zymo Research, Y1001M) in *Δbar1* deleted strains during 3 hours prior to imaging. G2/M arrest in metaphase was done through *P_MET3_-CDC20* (Paldi et al., 2020) depletion with 2 mM methionine added 2 hours before imaging. G2/M arrest in prometaphase was performed with nocodazole (Sigma, M1404), 20 ng/µl during 1h and 10 ng/µl for an additional hour. Smc5, Smc6 or Smc1 degradation was obtained by adding 1mM of auxin (Sigma, I3750-5G-A), dissolved in Ethanol 100%, 1 hour to the culture before imaging.

### Mass spectrometry

Purification of the endogenous yeast Smc5/6 complex was performed as described in (Adhikary and D’Amours, 2025) with minor modifications. A yeast culture (30L) was grown at 30 °C until it reached an OD_600_ of 0.7 – 1.0 and then further incubated at 18 °C for 16 hrs. Cells were collected and washed with H_2_O before being snap frozen. Cells were subsequently thawed on ice in NBE buffer (50 mM potassium phosphate buffer pH 8, 50 mM Tris-HCl pH 8, 500 mM NaCl, 10% glycerol, 0.5% Triton X-100, supplemented with 20 mM imidazole, 2 mM ß Mercapto-Ethanol, 2X protease inhibitors, pH 8). Snap frozen droplets of cell suspensions were lysed using a freezer mill. The resulting powder was then resuspended in NBE buffer and cleared by high-speed centrifugation. The cleared lysate was subjected to two consecutive rounds of Ni-NTA purification. The Ni-NTA columns were washed with 10 CV of NW buffer (50 mM potassium phosphate buffer pH 8, 50 mM Tris-HCl pH 8, 500 mM NaCl, 10% glycerol, 0.5% Triton X-100, supplemented with 60 mM imidazole, 2 mM ß Mercapto-Ethanol, and protease inhibitors, pH 8) and bond proteins were eluted with 5 CV of buffer NE (50 mM Tris HCl pH 8, 500 mM NaCl, 10% glycerol, 0.5% Tween-20, supplemented with 500 mM imidazole, 2 mM bME, and 1X protease inhibitors, pH 8). Eluted fractions were loaded on an SDS-polyacrylamide gel (4–12% Bis-Tris) in 1 x MOPS buffer and ran for 90 mins at 150 volts before being stained with Coomassie Blue. Peak fractions from both columns were combined and loaded on a 5mL Strep-Tactin XT column using an AKTA FPLC purification system. The column was washed with 10 CV of buffer SW (50 mM Tris HCl pH 8, 500 mM NaCl, 10% glycerol, 0.5% Tween-20, 2 mM ß Mercapto-Ethanol, 1X protease inhibitors, supplemented with 0.5% Triton X-100, pH 8) and eluted with 12 CV of STE buffer (150 mM NaCl, 10% glycerol in PBS 1X, supplemented with 50 mM Biotin, 2mM ß Mercapto-Ethanol, 1X protease inhibitors, pH 8). Peak elutions were snap frozen and stored at -80 °C

### MT binding and cell proliferation assays

The affinity of Smc5 hinge domain (residues 215 to 885) for MT polymers was monitored using a pelleting assay, as previously published (Laflamme et al., 2014). The proliferation phenotype of cells expressing wild-type and smc5 mutants was assessed by spotting 5-fold dilutions of yeast cultures on the surface of solid medium and monitoring yeast growth at the indicated temperatures after 2-4 days incubation at various temperatures (Laflamme et al., 2014).

### Co-immunoprecipitation (co-IP) of Smc5/6 complex subunits

The co-IP was performed as described in (Roy et al., 2024; under revision at eLife) with minor modifications. Yeast cultures (50mL) were grown at 30°C in YEPD medium to an optical density (OD_600_) of 0.7–0.8. Cells were harvested and lysed with glass beads in co-IP lysis buffer (50mM HEPES – KOH pH 7.5, 140mM NaCl, 1% Triton X-100, 0.1% sodium deoxycholate, 1mM EDTA pH 8.0, 1mM AEBSF, 10µM Pepstatin-A, 10µM E64). The purification resin (StrepTactin XT®) was washed with sterile water and PBS-BSA buffer (1x PBS [phosphate buffered saline], 1% BSA) just prior to the co-IP experiment. Cell lysates were cleared by centrifugation (30 minutes at 13,000rpm) and combined (1mL) with the washed resin for an overnight incubation (∼12h) at 4°C with gentle rotation (20rpm). The resin was subsequently washed in sequence with 1) co-IP lysis buffer, 2) high salt buffer (50mM HEPES – KOH pH 7.5, 360mM NaCl, 1% Triton X-100, 0.1% Sodium deoxycholate, 1mM EDTA pH 8.0) and 3) TE buffer (10 mM Tris-HCl pH 8.0, 0.1 mM EDTA). The purification resin was finally resuspended in 2X SDS-PAGE sample buffer and heated at 95°C for 5 minutes to elute purified proteins. The eluted proteins were separated on a SDS-polyacrylamide gel [4-12% Bis-Tris gel ran with 1x MOPS electrophoresis buffer (20mM MOPS [3-(N-Morpholino) propanesulfonic acid], 5mM of sodium acetate, 1mM of disodium EDTA [isodium ethylenediaminetetraacetate dihydrate]) for 2h at 120 volts, before being transferred to a membrane and analysed by immunoblotting using a 1:2,000 dilution of mouse anti-Strep antibody and 1:5,000 dilution of mouse anti-Myc (9E10) antibody (GeneTex 369661).

### Cell cycle time course experiment

To assess the kinetics of cell cycle progression, a time course experiment was performed as previously described (Ratsima et al., 2011). Specifically, yeast strains were grown in YPD medium at 30°C until they reached an OD_600_ of 0.3-0.4. Alpha factor was then added (5µg/mL) to the medium and, after 1h of growth, a second dose (2µg/mL) was added to ensure optimal arrest in G1. Once ≥ 90% of the cells were synchronized in G1, they were released at 30°C in fresh YEPD medium. Alpha factor (5µg/mL) was added 45 minutes after the initial release to prevent cell cycle re-entry after completion of the first mitosis. For each time point, 1.5ml of cells were collected and pelleted for 2 minutes at 3,000rpm. Cells were then fixed in 1mL of 70% ethanol overnight at 4°C. Cells were washed twice and resuspended in 0.1M KPO_4_ pH7.0 buffer. DAPI (4’,6- diamidino-2-phenylindole; 2µg/ml) was used to stain yeast nuclei. Separation of nuclei was manually counted in 100 cells per time-point (N=3). Fluorescent images were taken using a Nikon Eclipse Ti2 inverted microscope, with an 100x oil immersion objective.

### Flow cytometry

Yeast cells were grown in SC selective medium overnight at 30°C, diluted to O.D_600_ nm = 0.5 in 5 ml next morning for 1h30 at 30°C. If necessary, cells were arrested at G1 or G2/M. 10^7^ cells were centrifuged, fixed in 200 μl 70% ethanol and stored at 4°C for 48h. To perform FACS, cells were centrifuged at 10.000 rpm 5 minutes at 4°C, washed and resuspended in 1 ml of sodium citrate (50mM, pH7). 200 μl cells were transferred in polystyrene round-bottom tubes with 20 μL of RNaseA (10 mg/ml Roche, 10109169001) and incubated 1 h at 37°C. Cells were resuspended in 2 ml of sodium citrate (50mM, pH7) with 2.5 μM of SYTOX^TM^ Green nucleic acid stain (Sigma, S7020). Cells were stained 1h at 4°C in the dark for 1 hour. Cells were and analysed with the Attune NxT Flow cytometer.

### Microscopy

For microscopy, cells were grown as for FACS. Cells were plated on agarose patches (made of SC medium containing 2% agarose) and sealed using VaLaP (1/3 Vaseline, 1/3 Lanoline and 1/3 Paraffin). Cells were imaged using a widefield microscopy system Nikon Ti-E equipped with the Perfect Focus System (PFS) and a 60x oil immersion objective (Nikon, Plan APO). Andor Neo Scmos camera allowed a vision field of 276x233 μm with a pixel size of 108 nm. Fluorochromes were excited with 35% LED with a wavelength of 477 nm for GFP or 40% LED with a wavelength of 587 nm for mCherry.

### Dynamic analysis

Analyses of chromatin mobility were performed as described in Herbert et al., 2017. Briefly, GFP tracking was performed by video microscopy for 3 minutes with an interval time of 100 ms in two dimensions (x,y). A Fiji plugin was used to isolate and extract locus trajectory at each time point throughout the film. After drift correction and signal-to-noise ratio checked with MATLAB scripts, Mean square distances (MSDs) for each trajectory were calculated and concatenated. Power laws to the mean MSD of a population over time intervals between 0.1 and 10s were adapted with MATLAB script and graphs of averaged MSDs were plotted in linear or log/log.

For each single locus trajectory sampled at a time resolution *δt* for an overall time *T*, the MSD at time Δ*t* is calculated as the time-average over the 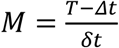 frames of the squared displacement performed by the locus between a running initial time *m δt* and the increased time *m δt* + Δ*t*:

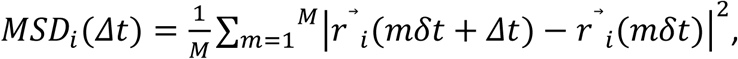

For an ensemble of *N* trajectories (cells), one gets an ensemble of *N MSD*_*i*_(Δ*t*) curves; the index *i* therefore refers to *i*-th the trajectory. Each single curve can then be fitted to a *y*(Δ*t*) = *D* ΔΔ*t*^*α*^function leading to a value of the exponent *α*. The distribution of the ensemble of the values of *α* obtained is shown in **Figure 1F**. In principle, the sub-diffusion coefficient *D* is also obtained by the same fitting procedure. However, we found that an equivalent, but more robust estimate of the sub-diffusion coefficient is obtained by the trend of the MSD over the very first 15 points, as 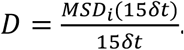 The *D* histograms of **Figure 1F** refer to this definition of the sub-diffusive coefficient.

The MSD can also be averaged over the ensemble of *N* trajectories (cells) as

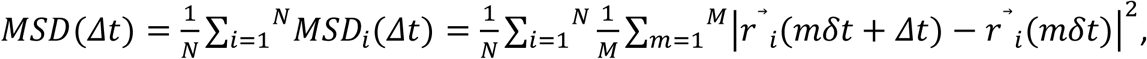

corresponding to the curves of **Figure 1E**.

### Distances analysis

The analysis of distances between two differently labelled dots (GFP and mCherry) was carried out either for intrachromosomal loci, or for a pericentromeric locus and the SPB. Acquisition of 3D z-stacks consist of 35 frames with z steps of 300 nm, illumination with two consecutive color channels for each plane with an exposure time of 100ms using a dual band filter set. First, using images from a strain dually labelled for the same locus, chromatic aberration was corrected using the plug-in “Descriptor based registration (2d/3d)”. The 3D z-stacks were then converted to 2D using a maximum intensity projection. The dots were then selected using Fiji and the (x,y) coordinates of the red and green loci calculated using a home-made script on Fiji, taking into account the Gaussian intensity of each fluorescent loci. Comparison of distances between loci was performed using Wilcoxon rank-sum tests.

### ChIP-qPCR

Yeast cells were grown either in YPD medium or SC-His Raff 2% overnight at 30°C. Next morning, cells were diluted to a O.D_600_ nm of 0,5 in 45 ml and cells were incubated at 30°C during 1h30 to reach the exponential phase. To check expression of Smc5 in the whole genome, cells were synchronized at G2/M phase using nocodazole (Sigma, M1404,). Nocodazole (cci. 2 mg/mL) was added two times during the time of synchronization, 20 ng/μL during a first 1h at 30°C and then 10 ng/μl for an additional hour at 30°C under agitation. Cell cycle arrest was verified by flow cytometry. In order to check the enrichment of Smc5 around the DSB, DSB was induced to the cell cultures by addition of 2% galactose during 6 hours at 30°C. 50 OD were recollected and treated with 1% formaldehyde for 12 minutes at room temperature with rotation. Crosslinking was quenched by using 2,25 mL of glycine 2,5 M for 5 minutes with rotation. Then, samples were harvested by centrifugation at 3000 rpm for 5 minutes and washed twice with cold TBS 1X.

Cells pellet was resuspended in 1,2 ml of lysis buffer (50 mM HEPES-KOH pH7.5, 140 mM NaCl, 1 mM EDTA, 1% Triton-X 100, 0.1% sodium deoxycholate) supplemented with 1 mM PMSF, phosphatase inhibitor, anti-protease (Roche, 04693132001) and 300 mg of glass beads (Sigma, G8772) and lysed by MagNA Lyser rotor (Roche, 03359093001). The lysate was obtained by centrifuging an eppendorf tube perforated with a sterile needle at 1000 rpm for 1 minute and recovering the extract in a new tube. The extract was transferred to a 1,5 ml falcon tube and sonicated with a PICO sonicator (15x: 30 sec ON, 90 sec OFF, high intensity) to obtain DNA with a range between 200 - 500 pb. After sonication cells were centrifuged for 10 minutes at 10.000 rpm at 4°C and 25 μL of the supernatant was saved at -80°C as INPUT. 250 μL of DNA after sonication were incubated with 1 μL of DNA carrier (ST0029, TakaRa Bio), 25 μL of BSA 5% (BP1600, Janssen Pharmacy) and 3,2 μg of anti-MYC antibody (Santa Cruz, Biotechnology, INC) overnight at 4°C. 30 ml of *Dynabeads™ Pan Mouse IgG* (Thermo Scientific, 11041) were added and incubated at 4°C for 4 hours. Beads were washed twice with lysis buffer, twice with lysis buffer + 0,36 M NaCl (50 mM HEPES-KOH pH7.5, 360 mM NaCl, 1 mM EDTA, 1% Triton-X 100, 0.1% sodium deoxycholate), twice with washing buffer (10 mM Tris-HCl pH8, 0.25 M LiCl, 0.5% NP40, 1 mM EDTA, 0.1% sodium deoxycholate) and once with elution buffer (50 mM Tris-HCl pH8, 10 mM EDTA). The supernatant was removed and the immunoprecipitated chromatin was then eluted by incubation in 250 μL TES buffer (50 Mm Tris-HCl, pH 8.0; 10 mM EDTA; 0,5% SDS) for 15 minutes at 65°C and the supernatant collected, termed IP. The INPUT was mixed with 120 μl of TES buffer. Both INPUT and IP were de-cross-linked at 65°C with agitation overnight. RNA was degraded by incubation with 2μL RNase A (10 mg/mL) for 1h at 37°C. Proteins were removed by incubation with 20μL of proteinase K (10 mg/mL) for 2h at 37°C. DNA was purified by a phenol/Chloroform extraction.

### Drop test

Overnight cultures were diluted to a O.D_600_=0.5 and cells were incubated at 30°C until they reached the exponential phase. Serial 1:5 dilutions were plated starting with a O.D.= 0.2 using the replication plate (R2383-1EA, Sigma) on YPD with or without 100nM hydroxyurea (Sigma H8627), 10 μM Camptotehcin (Sigma C9911) and 0.2 μM 4-Nitroquinoline 1-oxide (Sigma N8141). Plates were incubated at 30°C for 72 hours.

### Colony formation unit (CFU)

To quantify the colonies formed in the absence or presence of an I-*Sce*I induced DSB, cells were grown in -Ade -His + raffinose 2% liquid medium overnight at 30°C. After 24 hours, cells were counted in a Malassez chamber and ∼ 100 - 200 cells were spread on -Ade – His + glucose (control) or + galactose plates (DSB induction). Plates were incubated at 30°C and colonies were counted after 72h.

### Plasmid maintenance assay

Plasmid stability was assessed using the centromeric plasmid pRS415. WT and smc5-2KE mutant cells carrying the plasmid were grown overnight in YPD 200 cells were spread on YPD and replica plated on SC-Leu plates to determine the fraction of plasmid-retaining cells. Plasmid maintenance efficiency was calculated as the ratio of colony-forming units on SC–Leu versus YPD plates.

### Western blot

Electrophoretic protein separation was done using Bolt^TM^ 4-12% Bis-TrisPlus (Thermo Fisher NW04122BOX) in Mini gels tanks (Life Technology) and the electrophoresis buffer was NuPAGE^TM^ MOPS or MES SDS Running buffer (MOPS: Thermo Fisher NP0001; MES: Thermo Fisher, NP0002) for 24-26 minutes at 220 volts. Proteins were transferred in a BioTrace^TM^ nitrocellulose membrane (66485, Pall) in Mini PROTEAN Tetra System tanks (Bio Rad) for 90 minutes. at 100 volts. Membranes were blocked with 5% milk in Tween-TBS and then probed with 1/1000 Anti-MYC (Santa Cruz, Biotechnology), 1/1000 anti-HA (Sigma, H3663) or 1/500 anti-yH2A (Abcam, ab15083). Primary antibodies were recognized by secondary antibodies either anti-mouse or anti-rabbit IgG HRP-conjugated diluted 1/10000 (Thermo Fisher A11005

and A31556). Blots were developed using the ECL plus western blotting system (GE Healthcare) during 10-30 sec.

### qPCR

Cells were grown overnight in SC-Ade-His with 2% Raffinose at 30°C, diluted again in 2% Raffinose until they reached the mid-log phase and then, 2% galactose was added to the culture for I-*Sce*I induction. Cells were collected at 0, 2, 4 and 6 hours after galactose induction. DNA was extracted from each sample using phenol/chloroform extraction. For this purpose, cells were centrifuged 1 minute. at 13000 rpm, the pellet was mixed with 300 mg of glass beads, 200 μL of lysis solution (Triton X-100 2 %, SDS 1 %, NaCl 100 mM, Tris 10 mM pH 8, EDTA 1 mM) and 200 μL phenol-chloroform-isoamyl alcohol (Sigma, 77617). The mix was vortexed for 10 minutes at 4°C and centrifuged for 10 minutes. at 13000 rpm at 4°C. 140 μL of the supernatant was transferred into new tubes containing 500 μL ethanol 100% and DNA was precipitated during 1 hour at 80°C. The mix was centrifuged for 10 minutes. at 13000 rpm at 4°C. The pellet was washed with 500 μL of ethanol 70 % and dried at room temperature resuspended in 40 μL of TE containing 20 μg/μL of RNase (Roche, 10109169001) and incubated 30 minutes. at 42°C. DNA concentrations were quantified using nanodrop.

qPCRs were prepared in a 360-wells plate. All samples were brought to a concentration of 5 ng/μL. In each well, 4μL of the sample were mixed with 4 μL of Power Track SYBR^TM^ Green master mix (Thermo Fisher A46109) and 0,3 μL of forward and reverses oligonucleotides (final concentration 2.10^-5^ M). Oligonucleotides are listed in **Table 2**. I-*Sce*I oligonucleotides generated a 187bp fragment that flanks the I-*Sce*I cutting site. ACT1 was used as a control to normalize the samples. The standard curve was performed on actin (control) and I-*Sce*I cutting site (before cut) to evaluate the efficiency of the couple of primers used. Triplicates from each sample were performed and four 1:5 dilutions were performed for the standard curve.

**Table 2:**
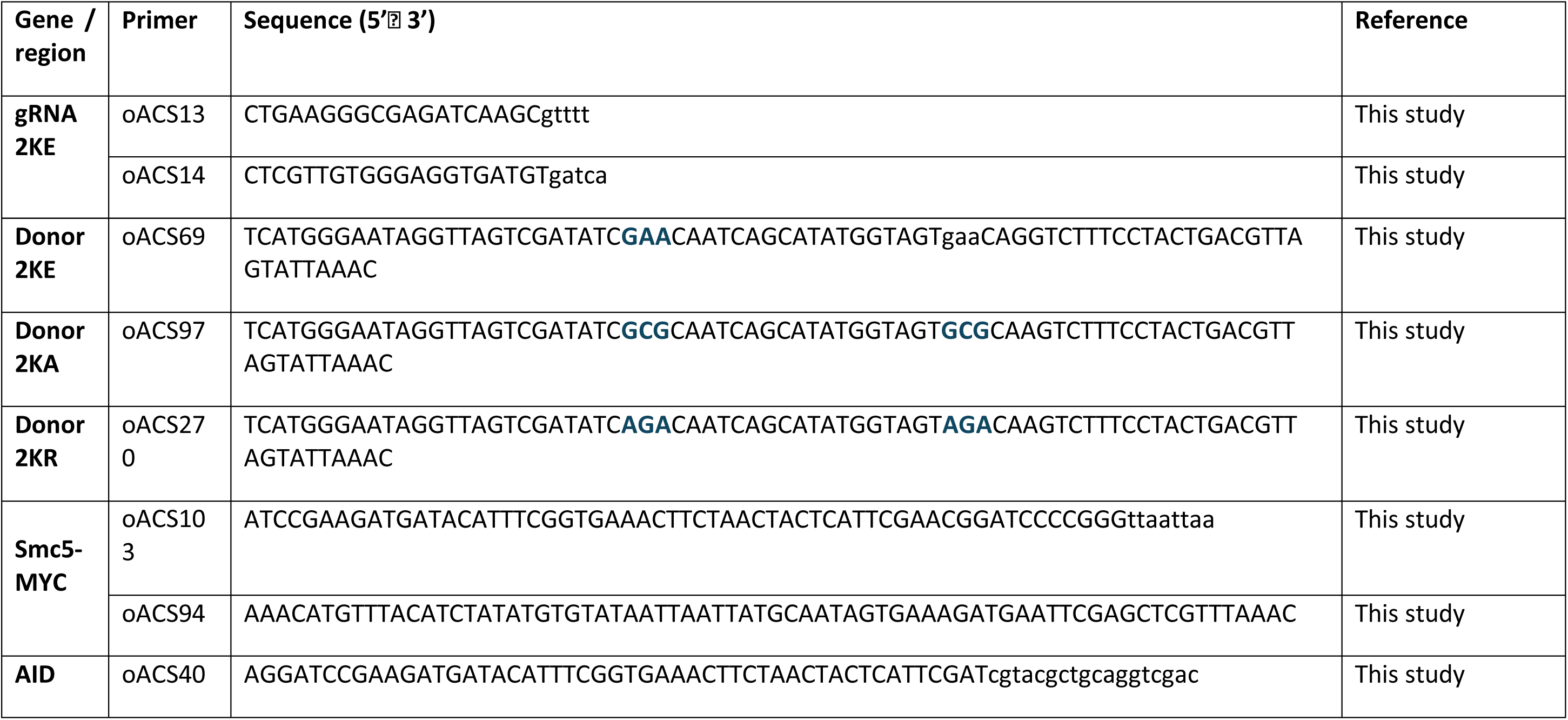
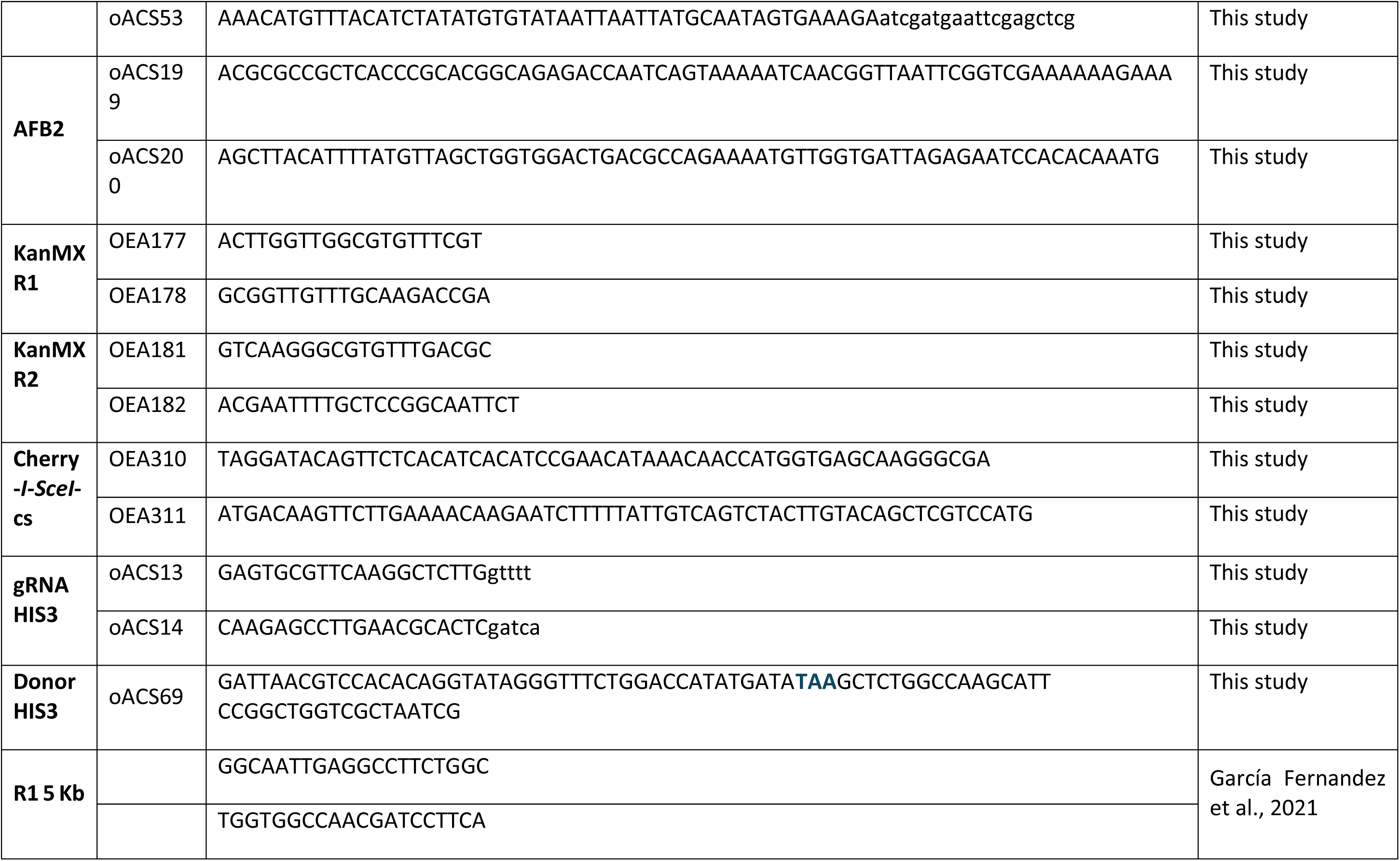
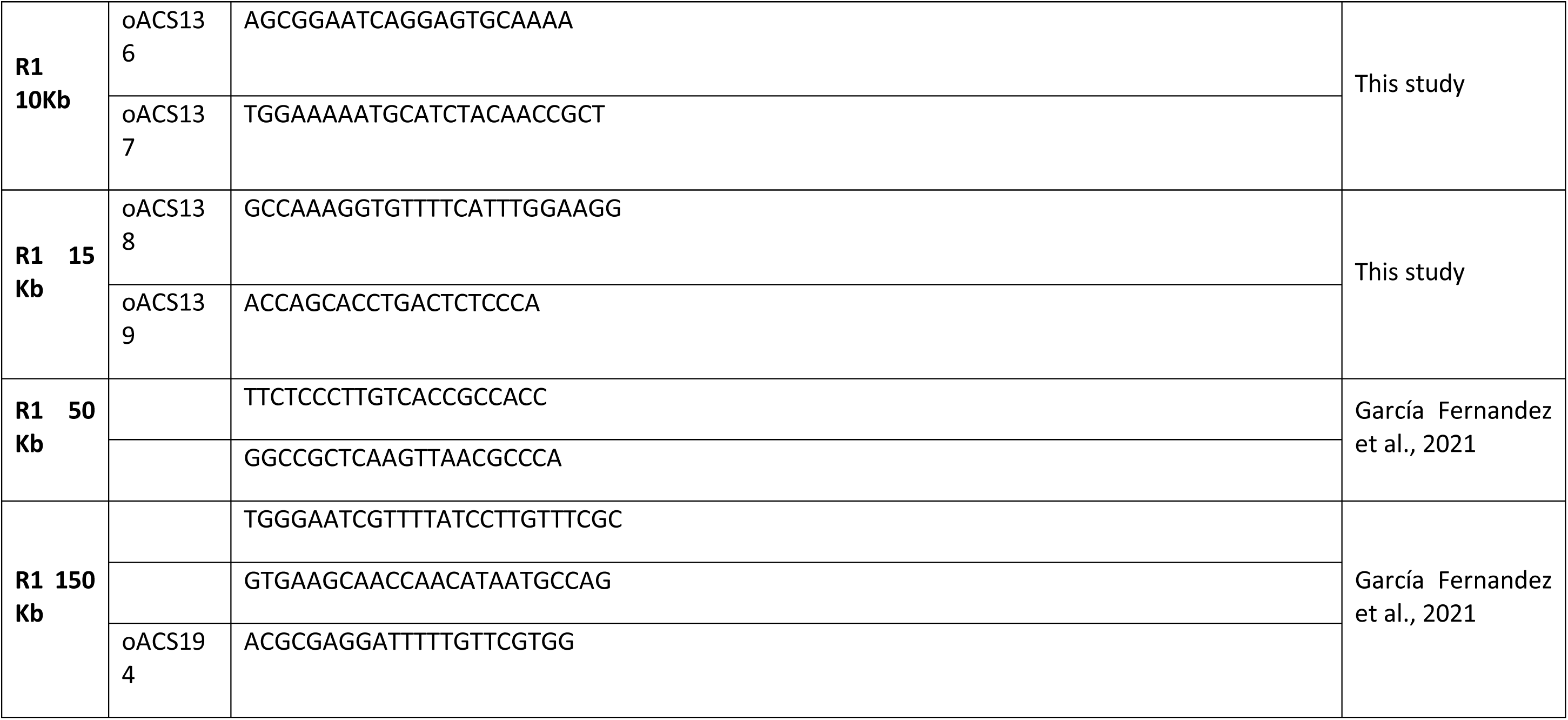
primers used in this study.

**Table 3:** plasmids used in this study.

| Name | Marker | Features | Origin |
| --- | --- | --- | --- |
| **pEF562** | KANMX | Plasmid carrying Cas9 and a gRNA scaffold. gRNAs were introduced by digesting this plasmid with BpII. | 87 |
| **pEF590** | KANMX | Plasmid coming from the pEF562 carrying the gRNA to cut close to the SMC5-K631. | This study |
| **pEF601** | URA3 | Plasmid carrying pFA6a-13MYC-URA3 used to introduce 13x MYC at the C-terminus of *SMC5* | P. Lesage |
| **pEF222** | LEU2 | Plasmid carrying pTEF-tTEF-LEU2-LacO, used to integrate the LacO in the place of a KanMX | Therizols et al., 2010 |
| **pEF569** | HIS3 | Plasmid carrying LacI-GFP-HIS3, which was linearized with *NheI* to introduce LacI-GFP into *HIS3* |  |
| **pEF604** | HIS3 | Plasmid carrying the AFB2, which was linearized with *NheI* to introduce AFB2-URA3 into *URA3*. | T. Teixeira |
| **pEF608** | HIS3 | Plasmid coming from the pJH2970 (Addgene, 100954) carrying the gRNA to cut close to the Smc5-K631. | This study |
| **pB07** | HIS3 | Plasmid carrying the I-SceI enzyme under GAL1-10 promoter | García-Fernandez et al 2022 |
| **pEF573** | ADE2 | Centromeric plasmid carrying the mCherry donor sequence, without the promoter and terminator. | García-Fernandez et al 2022 |

Plate sampling was distributed by the epMotion 5070 robot (Eppendorf), qPCR was performed with QuanStudio5 qPCR machine (Thermo Fisher) under the following conditions: pre-incubation (95°C – 5 min – 4.4 °C/min), amplification (45 cycles; 95°C – 10 s - 4.4°C/min; 58°C – 10 s - 2.2 °C/min; 72°C – 20 s – 4,4 °C/min), melting curve (95°C – 5 s - 4.4 °C/min; 60°C – 1 min – 2.2 °C/min; 97°C – continuous – 0.06 °C/min; 10 acquisitions/°C) and cooling (40°C– 30 s - 2.2 °C/ min.). qPCR was analyzed with Thermo Fisher cloud and normalized by the Livak method (2^ΔΔCt^) (Livak and Schmittgen, 2001).

### Polymer model

The polymer model was identical to the one used in Liu et al. 2023 (Liu et al., 2023) which was based on the model of Arbona et al. 2017 (Arbona et al., 2017) but made in Polychrom (https://github.com/open2c/polychrom/tree/master), a python wrapper for polymer simulations using openMM (Eastman et al., 2017). The model consisted of a diploid yeast nucleus where each monomer represents 750 bp and is 15 nm in radius. The monomers were connected with harmonic bonds with a spring constant parameterized by a wiggle-distance w such that bond extensions of w lead to bond energies of 1kT

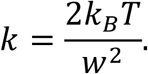

From this follows that, in the absence of other forces, the wiggle distance will be equal to 2 times the bond standard deviation.

The beads were allowed to interact through the soft “polynomial_repulsive” potential. Stiffness was introduced through a harmonic force on the bond angle which favored alignment. The spring constant was chosen to yield a persistence length of 70 nm which was found optimal by Arbona et al. The nuclear envelope was modeled by a spherical confinement with a 1µm radius, and all telomeres were attached to the wall using strong harmonic bonds which allowed movement along the periphery but no radial motion. All centromeres were attached to the SPB using harmonic bonds with a 400 nm rest length and a high spring constant, allowing negligible fluctuations in length. The nucleolus was modeled by inserting 112,500 bp at the rDNA region where the size of monomers and bonds were increased by a factor 14. Long simulations were run to collect a set of starting configurations from which the starting conformation for short-time simulations were then randomly chosen upon initialization. The conversion factor between simulation time and experimental time was chosen to match the MSD of the 11kb loci in the WT condition.

### Alphafold3 model

To predict potential interactions between the Smc5 hinge domain and candidate binding partners, the Smc5 hinge domain (residues 450–645) was modeled in complex with full-length (Tub1-Tub2), actin (Act1), or Shs1 subunit of the septin complex from *Saccharomyces cerevisiae*. Protein sequences were provided in FASTA format, and structural predictions were performed using default AlphaFold3 parameters. The confidence of the predicted interactions was evaluated using the interface predicted TM-score (iPTM), which reflects the likelihood of a specific protein-protein interaction.

## Acknowledgements

We thank A. Marston, P. Hieter and T. Teixeira for sharing strains and plasmids. We also thank E. di Santo and C. Marquez for their participation in strains constructions. Our special thanks go to J. Reguera for his expertise on alpha-fold and his help on protein interaction modeling. Our acknowledgements go to S. Grosse-Holz, X. Zhao and J-M. Victor for enlightening discussions, P. Lesage, F. Garcia-Fernandez and S. Mihic for their constructive reading of the manuscript. P. Therizols and the members of the laboratory are thanked for the many lively discussions on this project and the GDR Architecture Dynamique du noyau et des génomes (ADN&G) for the valuable exchanges with its community. This work was supported by an FRM grant to EF (EQU202203014635) and CIHR grant to DD (FDN–167265). EF is also supported by PLBIO-INCA (PLBIO20-31) and DD by a Canada Research Chair in Chromatin Dynamics & Genome Architecture (CRC-2017-00064). ACS was supported by FRM (ECO202006011576) and ARC (ARCDOC42023010006078) fellowships.

---

**Figure Supplement 1.**
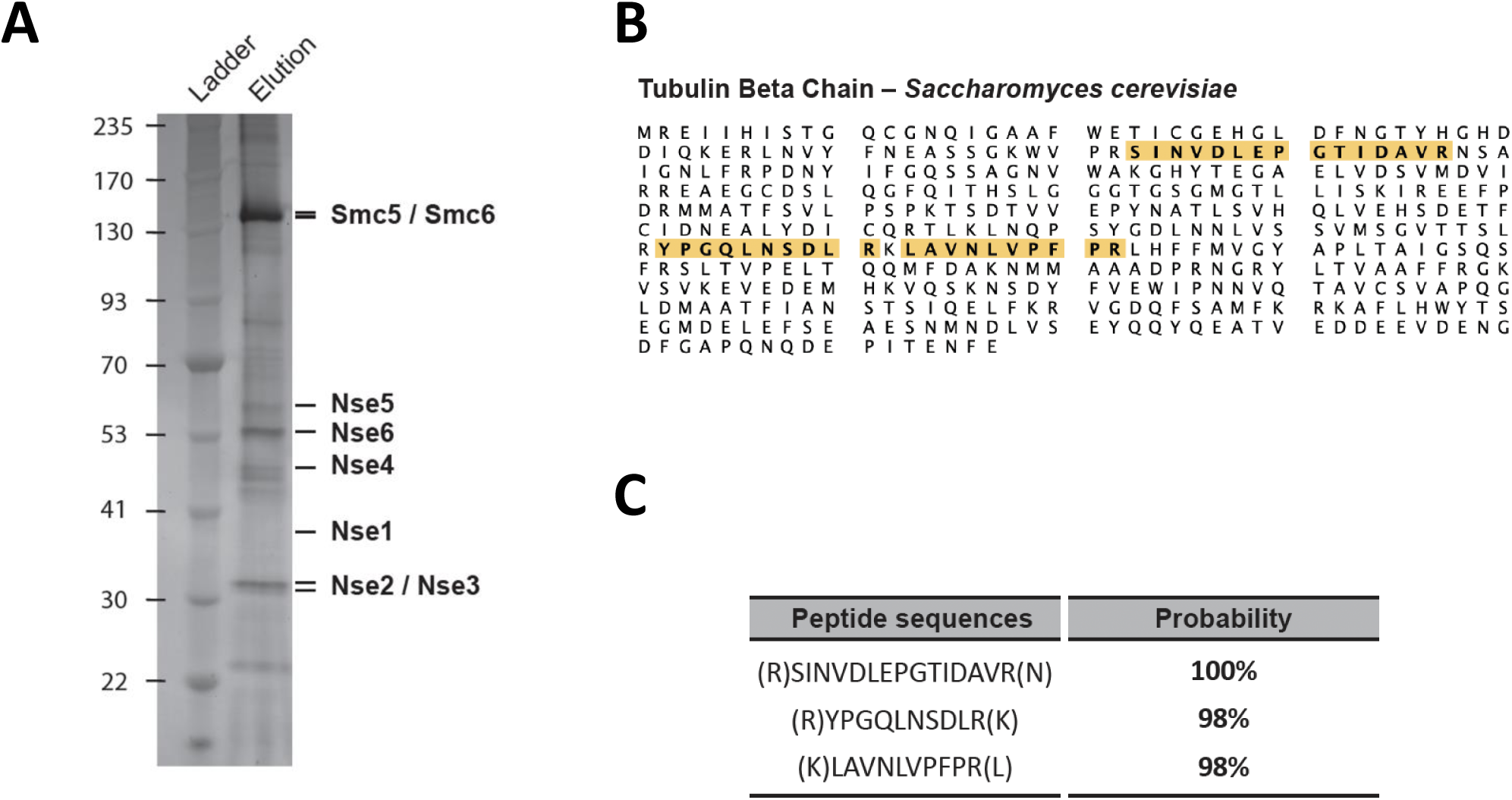
Mass spectrometry analysis of proteins co-purifying with the yeast Smc5/6 complex. The Smc5/6 complex was isolated from crude extracts using Ni-NTA and Strep-Tactin affinity chromatography and purified proteins were subjected to mass spectrometry analysis. **(A)** Silver-stained gel showing the 8 subunits of the yeast Smc5/6 complex obtained after consecutive rounds of Ni-NTA and Strep-Tactin affinity chromatography. The expected positions of Smc5/6 subunits are marked next to the gel. **(B)** Amino acid sequence of the beta chain of yeast tubulin (Tub2). High probability peptide sequences identified in the mass spectrometry analysis are highlighted in yellow. **(C)** Tub2 peptides detected by mass spectrometry and their probability of identification.

**Figure Supplement 2.**
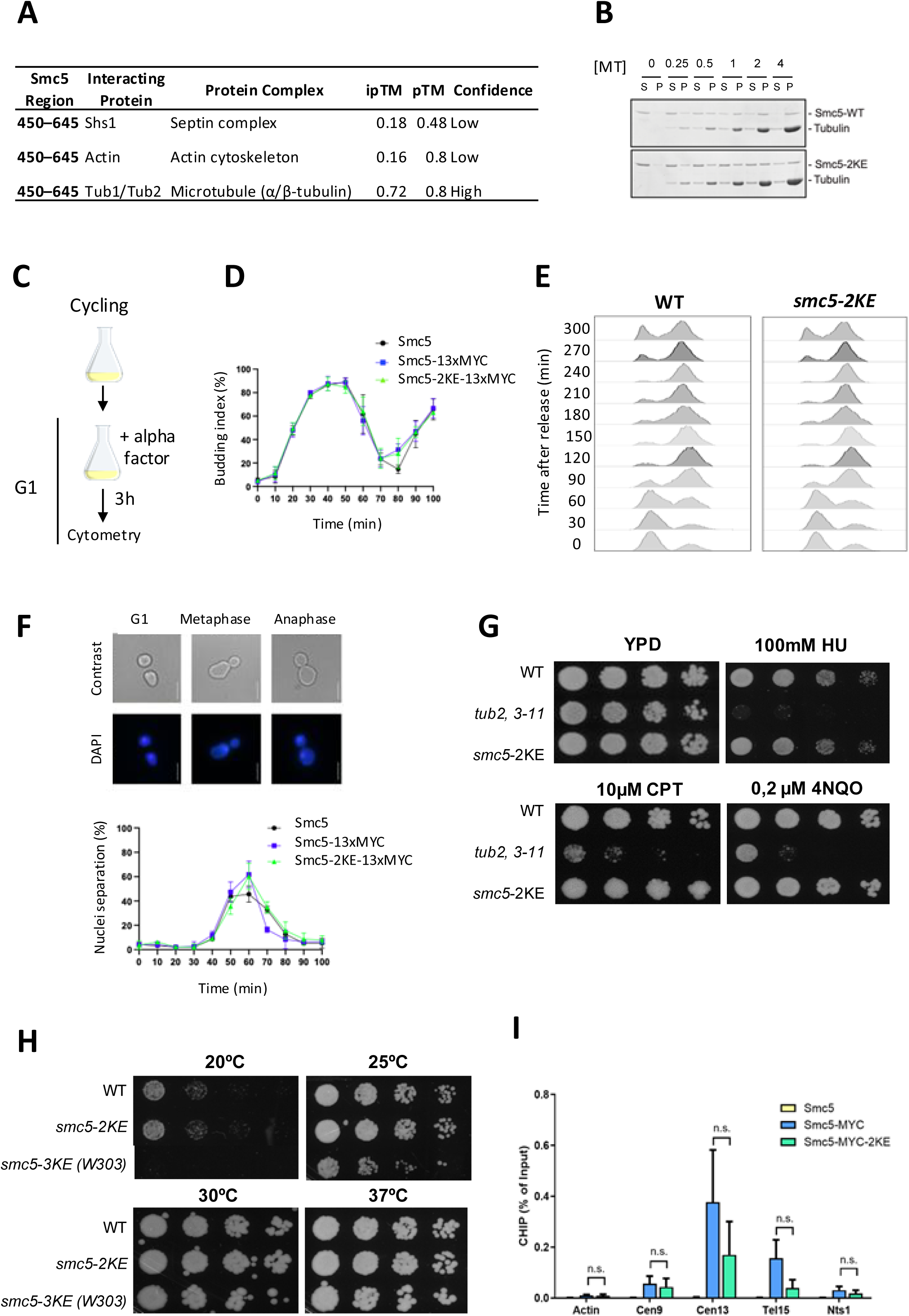
Smc5-2KE mutation does not affect cell growth nor cell cycle. **(A)** Table showing AlphaFold3 predictions of interactions between the Smc5 hinge domain (residues 450–645) and candidate proteins. Only the Tub1/Tub2 complex showed a high-confidence interaction (ipTM = 0.72), while Shs1 displayed low ipTM scores, indicating weak or no predicted interaction**. (B)** Microtubule polymer binding assay. Coomassie blue-stained gels showing the binding of the hinge region of Smc5 (residues 215–885) to the microtubule (MT, µM) polymer fraction (pellet; P) after centrifugation. Note how Smc5 hinge domain progressively transferred from the depolymerized tubulin fraction (supernatant; S) to the polymer fraction (P) as the MT concentration increased and eventually bound all available Smc5 protein (top gel). In contrast, the Smc5-2KE mutant (lower gel) showed strongly reduced affinity for MT polymers, as evidenced by the substantial fraction of the hinge domain remaining in the supernatant fraction at the highest concentration of MT shown in the gel (4 µM). **(C)** Experimental protocol used to synchronize cells in G1 using alpha factor. **(D)** Percentage of budded and late anaphase cells at each time point, respectively. **(E)** Cell synchronization monitored by flow cytometry. **(F)** Images showing examples of G1 (**left**), metaphase (**middle**) and anaphase (**right**) cell morphologies monitored in yeast cultures expressing Smc5 o*r* smc5-2KE. Prior to imaging, cells were fixed, washed and stained with DAPI to enable nuclei visualization. Scale bars represent 5µm. **(G-H)** Growth phenotypes on solid medium **(G)** in the presence of DNA-damaging agents — 100 mM hydroxyurea (HU), 10 μM camptothecin (CPT), or 4-nitroquinoline-1-oxide (4NQO) and **(H)** at different temperatures. 1/10 serial dilutions were spotted and photographed after 48h **(I)** Chromosomal association of Smc5-MYC at selected loci in the indicated strains, as determined by ChIP-qPCR. Cells were arrested in G2/M prior to sample collection. The Y-axis represents the amount of DNA in the ChIP fraction relative to input. Primers were designed to amplify the centromeres of chromosomes 9 (Cen9) and 13 (Cen13), the *rDNA* non-transcribed spacer 1 (Nts1), telomere 15 (Tel15). *ACT1* gene is used as a control. Each bar shows the mean of three independent experiments, with standard deviations indicated. Statistical significance was assessed using a two-tailed Student’s t-test. P-values are denoted as follows: n.s. = not significant (p > 0.05).

**Figure Supplement 3.**
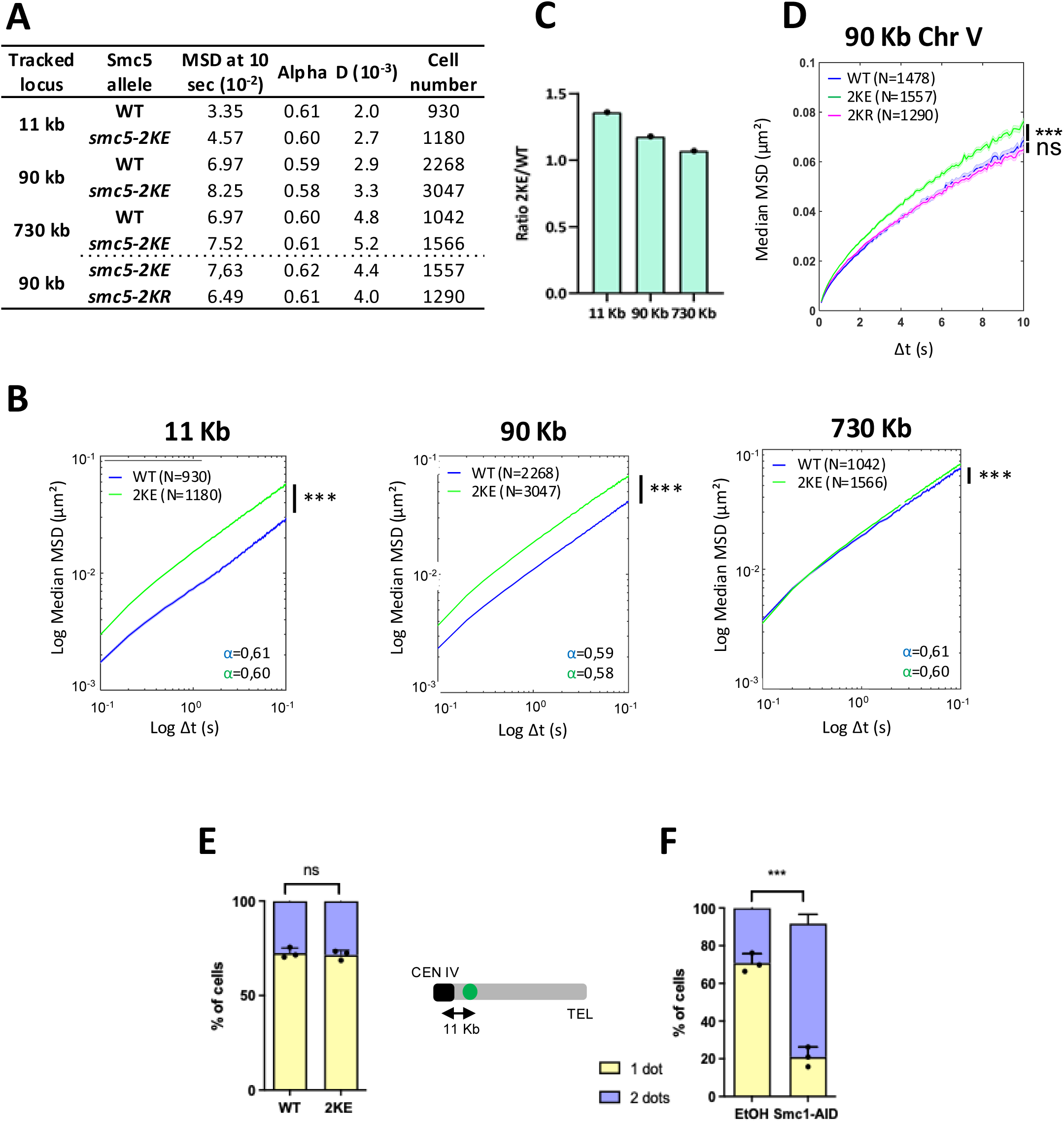
The negative charges of K624E and K631E, and not post-translational modifications, lead to increased chromatin mobility along the chromosomal arm in smc5-2KE with no effect on chromatin cohesion. **(A)** Table summarizing the different conditions with values extracted from the MSD: MSD at 10 s, the anomalous exponent (alpha), the sub-diffusive coefficient (D) and the number of cells analyzed. **(B)** MSDs as a function of increasing time intervals on a log-log scale in the WT (blue) and in the *smc5*-2KE mutant (green) for a locus at different positions of chromosome IV (11kb, 90 kb and 730 kb, from left to right). Wilcoxon rank-sum test results between distributions, with the p-value. * (p<0.05), *** (p<0.0001). **(C)** Ratio of MSD at 10 s between *smc5-2KE* versus WT at different positions of the chromosome IV. **(D)** MSDs as function of time in the WT, *smc5-2KR* and *smc5-2KE* mutants. Wilcoxon rank-sum test results between distributions, with the p-value. ns (p>0.05) and *** (p<0.0001). **(E-F)** Cohesion status was assessed based on the number of GFP foci separation of tetO/TetR-GFP marker at ∼11 kb from the centromere of chromosome IV in metaphase. Cells were synchronized by Cdc20 depletion. **(E)** WT Smc5 or the *smc5-2KE* mutated cells, and **(F)** cells in which Smc1 was tagged with an auxin-inducible degron. Cells with a single GFP (yellow) dot were scored as having cohesive sister chromatids, while cells with two GFP foci (purple) were scored as having defective cohesion. For each biological replicate, 200 cells were scored. Data show means of three replicates, with error bars representing ± sd; unpaired two-tailed t-test was used to analyze differences between groups, with “ns” for not significant (P>0.05) and * for P<0.01.

**Figure Supplement 4.**
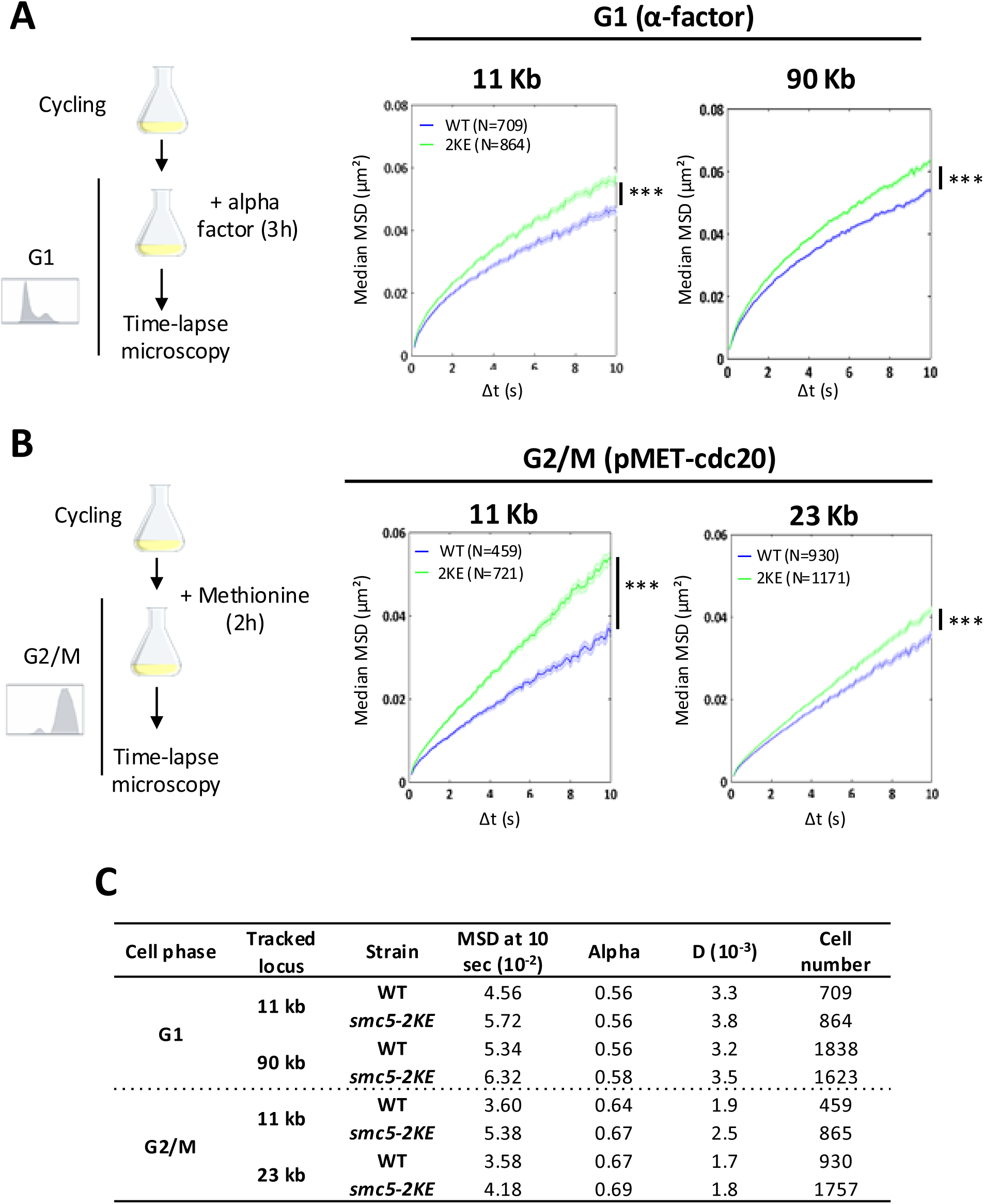
Smc5-2KE mutation increases chromatin mobility throughout the cell cycle. **(A)** Experimental protocol used to synchronize cells in G1 with 50 ng/ml of alpha factor (**left**). WT and *smc5*-2KE MSDs at 11 and 90 Kb at G1. Wilcoxon rank-sum test results between distributions, with the p-value *** (p<0.0001) (**right**). **(B)** Experimental protocol used to synchronize in G2/M cells carrying the *P_MET3_-CDC20* with 2 mM of methionine (**left**). WT and *smc5*-*2KE* MSDs at 11 and 23 Kb at G2/M phase. Wilcoxon rank-sum test results between distributions, with the p-value *** (p<0.0001) (**right**). **(C)** Table summarizing the different conditions with values extracted from the MSD: MSD at 10 sec, the alpha, the sub-diffusive coefficient (D) and the number of cells analyzed.

**Figure Supplement 5.**
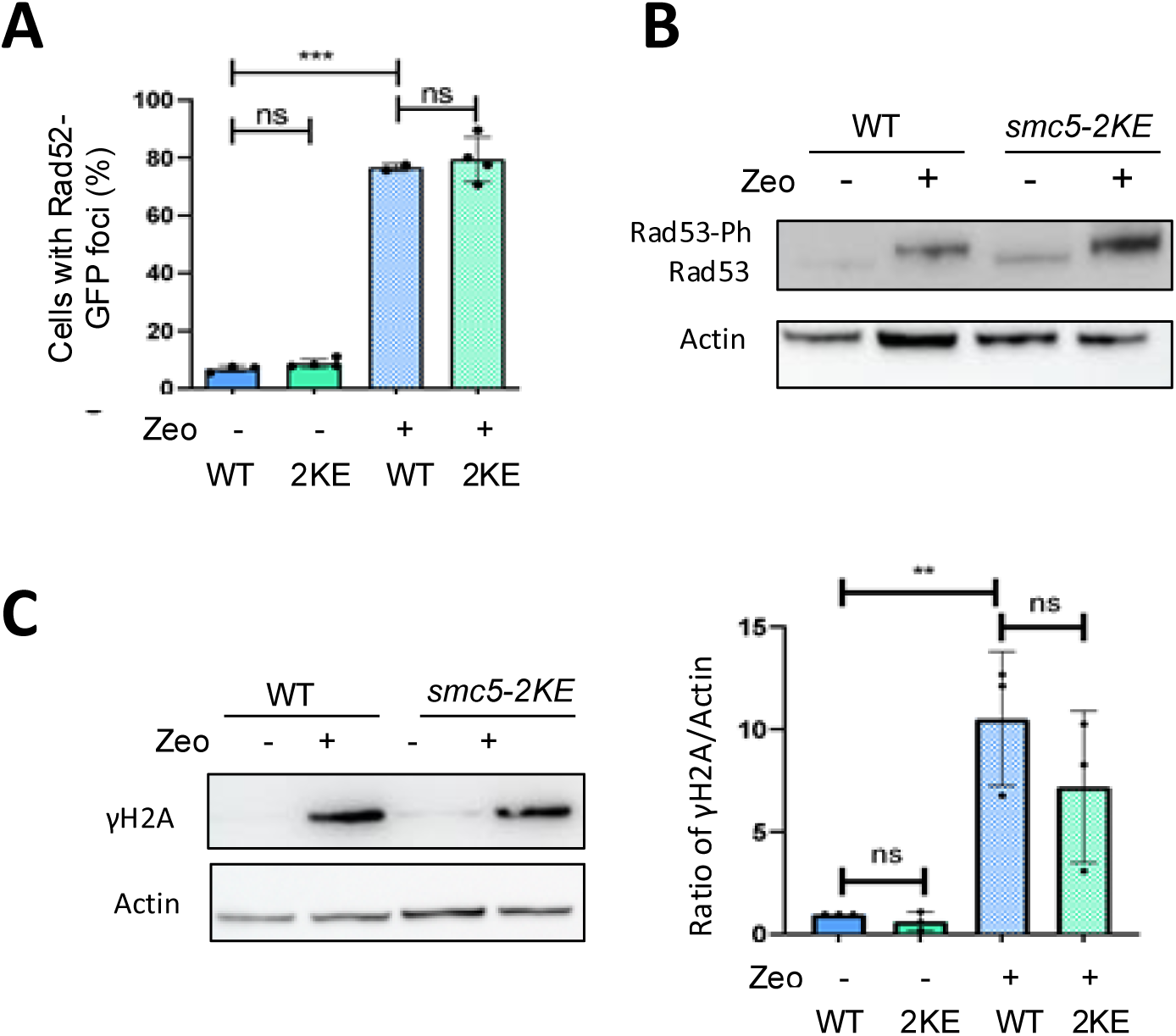
The *smc5-2KE* mutant does not induce spontaneous DNA damage. **(A)** Rad52-foci (protein involved in HR) in WT and *smc5-2KE* mutant (2KE). Three independent experiments were conducted with the number of cells counted as follows: N_WT_=608, N_2KE_=1388, N_WT ZEO_=650, N_2KE ZEO_=509. T-test was used to analyze differences between groups, with “ns” for not significant and *** for P<0.0001. **(B)** Representative results from immunoblotting showing the phosphorylation (Ph) status of Rad53 in WT, *smc5-2KE* cells and in response to Zeocin treatment (zeo). Actin was used as a loading control. Blots are representative of three experiments. **(C)** Representative results from immunoblotting showing the phosphorylation of H2A (γH2A) in WT, *smc5-2KE* cells and in response to Zeocin treatment. Actin was used as a loading control. Blots are representative of three experiments (**left**). Protein levels are represented as mean ratio values quantified from protein γH2A versus actin (**right**).

**Figure Supplement 6.**
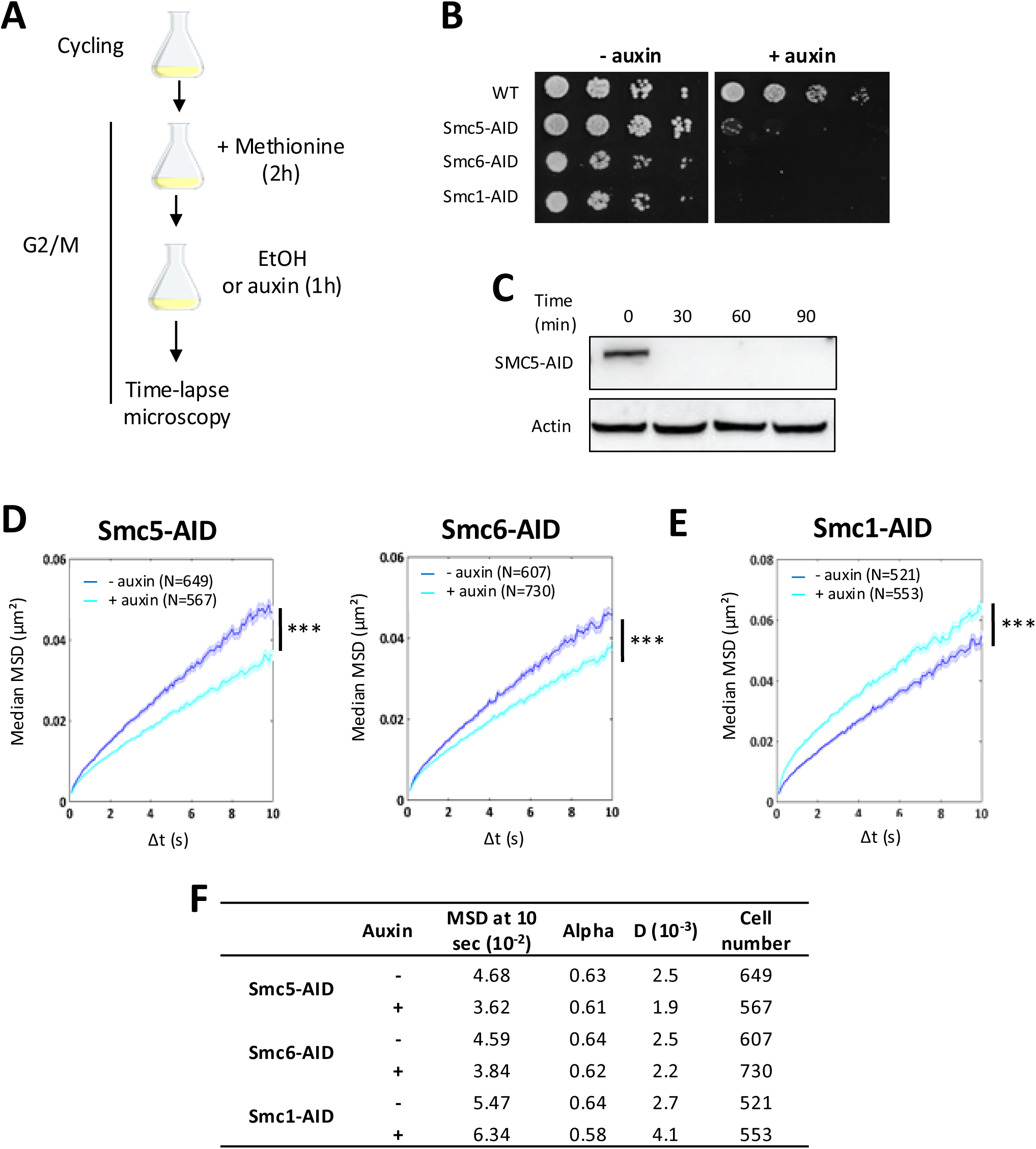
The Smc5 or Smc6 degradation decreases chromatin dynamics. **(A)** Experimental protocol used to synchronize cells in metaphase with 2 mM of methionine cells carrying the *P_MET3_-CDC20*, followed by degradation of SMCs with 1mM auxin for 1 hour prior to imaging. Auxin being diluted in ethanol (EtOH), controls with EtOH only were performed **(B)** Growth phenotypes on solid medium with or without Auxin in WT, Smc5-AID, Smc6-AID and Smc1-AID **(C)** Representative results from immunoblotting showing the degradation of the Smc5 upon 30 minutes of treatment with 1 mM auxin. Actin was used as a loading control. Blots are representative of three experiments. **(D-E)** MSDs as a function of increasing time intervals in the absence of auxin (dark blue) and in the presence of Auxin (light blue) in a locus at 11kb from the CEN4 in **(D)** Smc5-AID, Smc6-AID and **(E)** Smc1-AID. Wilcoxon rank-sum test results between distributions, with the p-value *** (p<0.0001). **(F)** Table summarizing the different conditions with values extracted from the MSD: MSD at 10 sec, the anomalous exponent (alpha), the sub-diffusive coefficient (D) and the number of cells analyzed.

**Figure Supplement 7.**
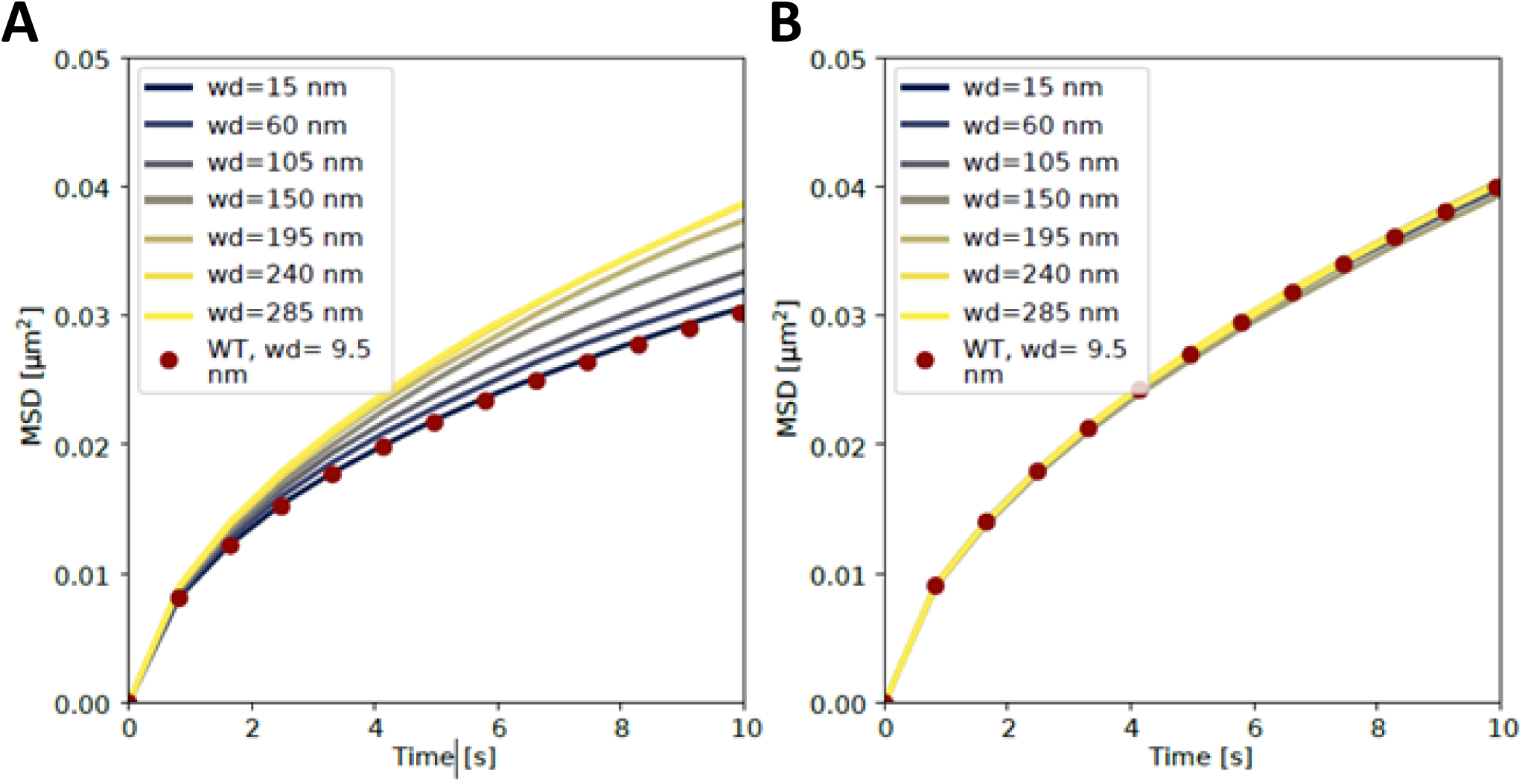
Weakening the microtubule tether to the kinetochore gradually abolishes position-dependent mobility: Sweep of the wiggle distance (wd) of the bond connecting the centromere to the spindle pole body (See material and methods for a mathematical definition of wd. In the absence of other forces it is √2 times the bond standard deviation). **(A)** MSD of loci 11 kb from the centromere. **(B)** MSD of loci 90 kb from the centromere. Candidate loci are selected as in Figure 4 and wild type (WT) refers to the simulation in Figure 4A.

**Figure Supplement 8.**
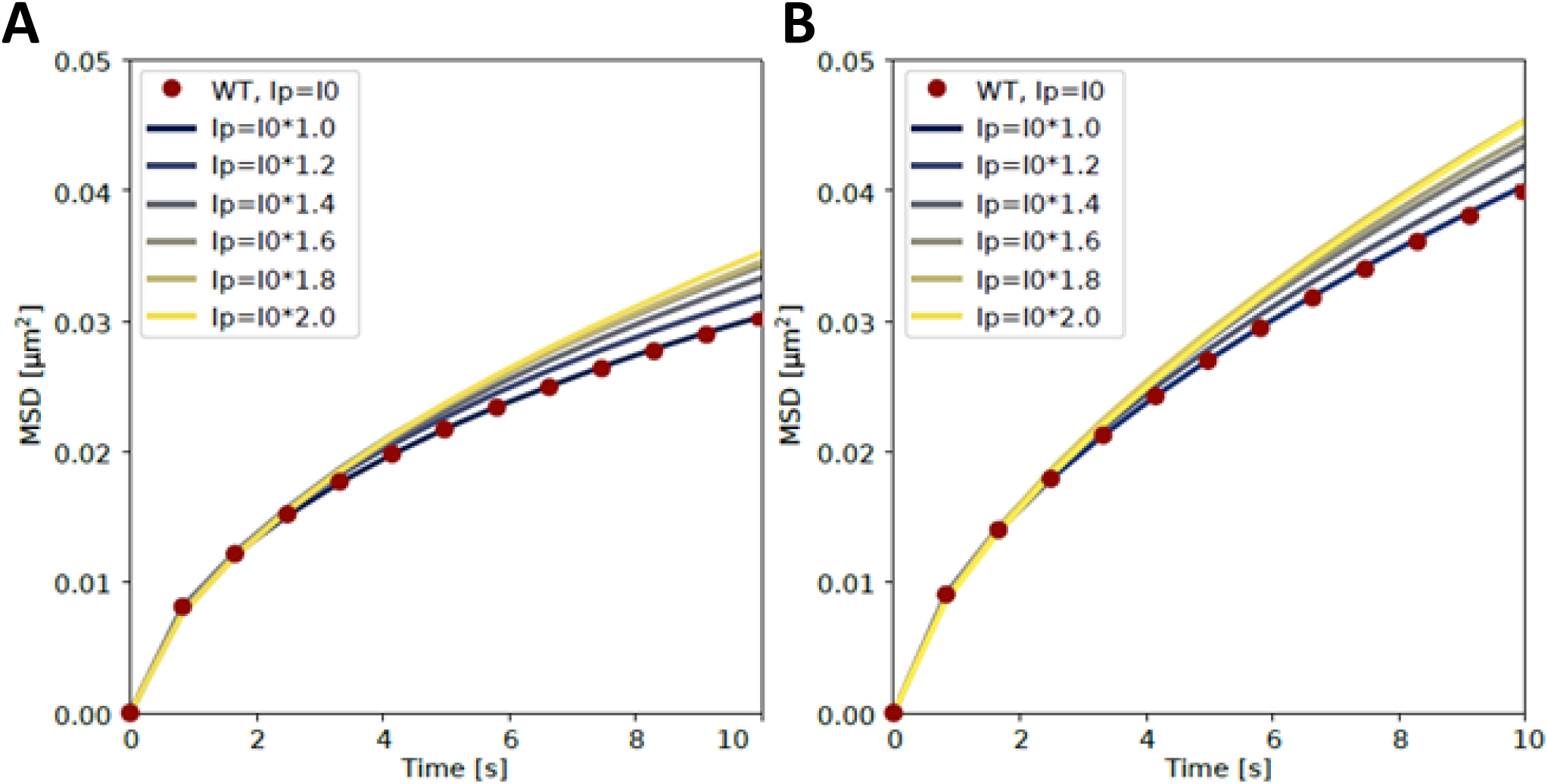
**Increasing chromatin stiffness increases mobility globally**: Sweep of chromatin stiffness by increasing the stiffness of the bond angle potential by up to a factor two. **(A)** Loci 11kb from the centromere **(B)** Loci 90kb from the centromere. Candidate loci are selected as in Figure 4 and wild type (WT) refers to the simulation in Figure 4A.

**Figure Supplement 9.**
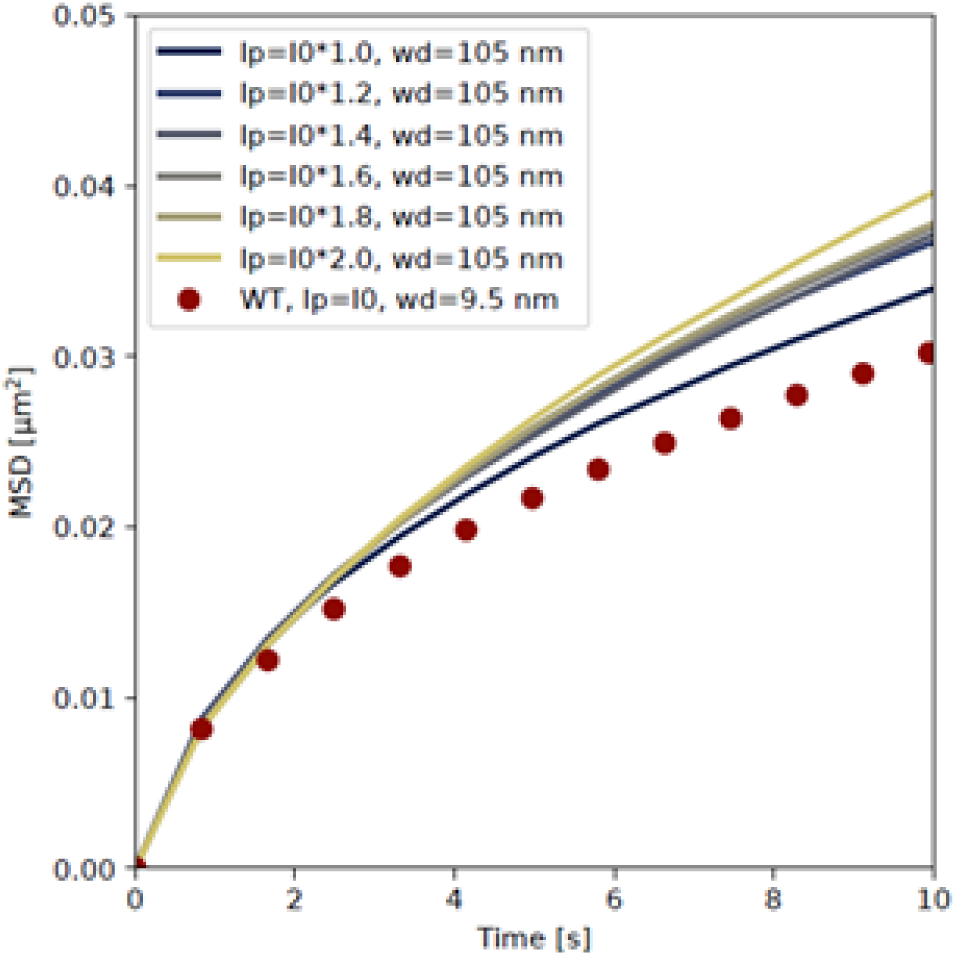
Combining chromatin stiffness and loosening the kinetochore tether further increases mobility in the pericentromeric region: Sweep of chromatin stiffness as in Figure supplement 7 but with a wiggle distance (wd) of 105 nm on the bond connecting the centromere and the spindle pole body. Candidate loci are 11 kb from the centromere and selected as in Figure 4. Wild type (WT) refers to the simulation in Figure 4A.

**Figure Supplement 10.**
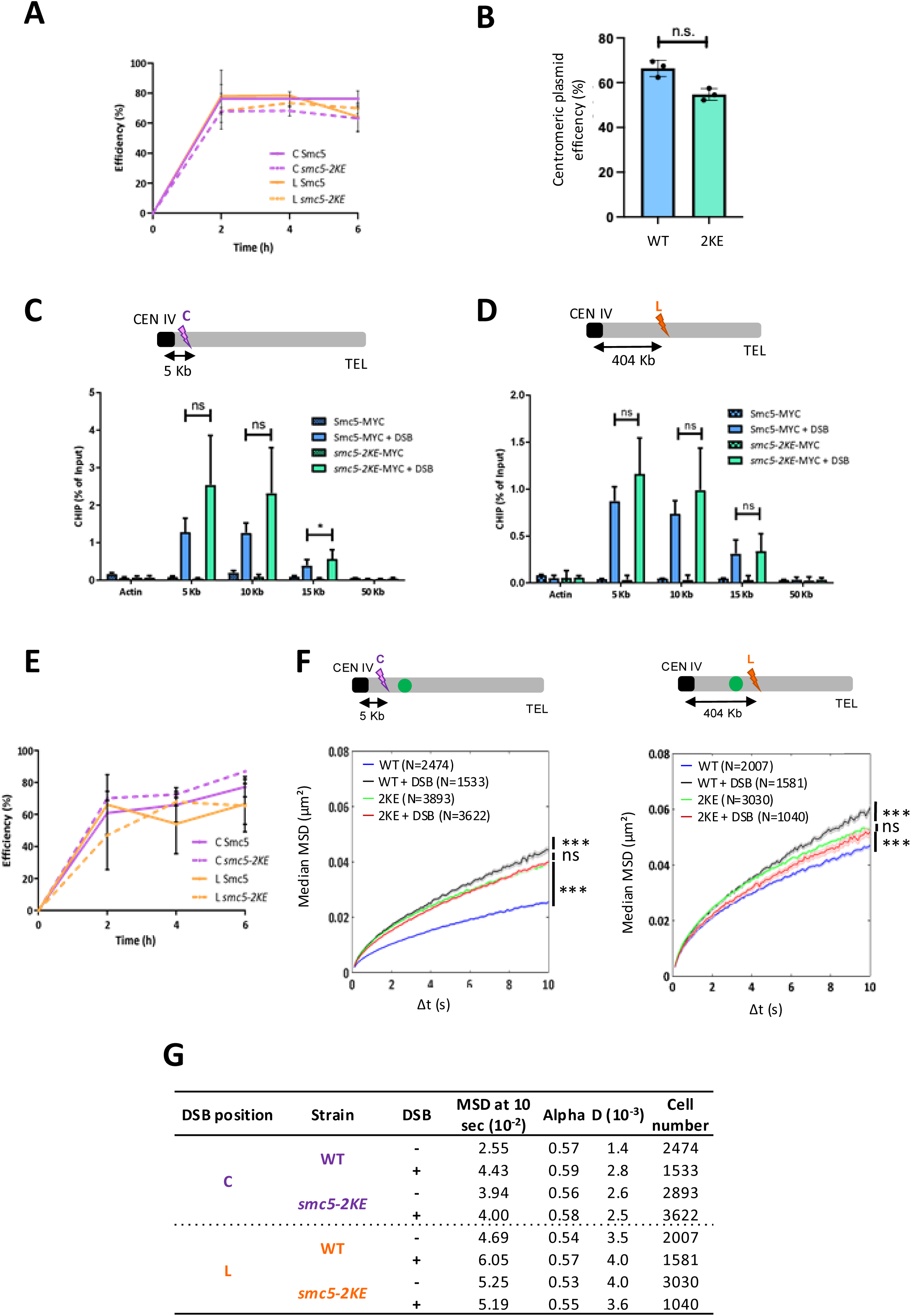
Smc5-2KE mutation affects HR at pericentromeres without affecting Smc5/6 chromatin loading. **(A)** I-*Sce*I efficiency (%) after 0h, 2h, 4h and 6h after inducing the I-*Sce*I with galactose in Smc5 WT and *smc5-2KE* for strains with a DSB at centromeric (C) or luminal (L) positions. qPCR was done using primers flanking the I-*Sce*I cutting site and actin was used as a loading control. The error bars represent the standard deviation of three independent experiments**. (B)** Proportion of cells retaining a centromeric plasmid after growth in nonselective conditions. Mann-Whitney test was used to analyze differences between groups, “ns” for not significant. **(C,D)** Chromosomal association of Smc5-MYC and Smc5-2KE-MYC at selected chromosomal positions as determined by ChIP-qPCR. Chromosomal positions from the DSB are indicated in the X-axis. Unpaired two-tailed t-test was used to analyze differences between groups, with “ns” for not significant (P>0.05) and * for P<0.01. **(E)** I-*Sce*I efficiency (%) of the Smc5 and *smc5-2KE* for C and L strains, used for local mobility **(F)** MSD as a function of increasing time intervals in the WT and *smc5*-2KE mutant after inducing a single DSB either at position C or L. Wilcoxon rank-sum test results between distributions, with the p-value “ns” non-significant, *** (p<0.0001). **(G)** Table summarizing the different conditions with values extracted from the MSD for the fourth strains used in local mobility: C WT, C *smc5-2KE,* L WT and L *smc5-2KE*: MSD at 10 sec, the anomalous exponent (alpha), the sub-diffusive coefficient (D) and the number of cells analyzed.

